# Autosomal Allelic Inactivation: Variable Replication and Dosage Sensitivity

**DOI:** 10.1101/2025.08.13.670061

**Authors:** Michael B. Heskett, Athanasios E. Vouzas, Brian Johnstone, Krister P. Freese, Phillip A. Yates, Philip F. Copenhaver, Paul T. Spellman, David M. Gilbert, Mathew J. Thayer

## Abstract

Autosomal monoallelic gene expression and asynchronous replication between alleles are established features of imprinted genes and genes regulated by allelic exclusion. Inactivation/Stability Centers (I/SCs) are recently described autosomal loci that exhibit epigenetic regulation of allelic expression and replication timing, with differences that can be comparable to those observed between the active and inactive X chromosomes^1^. Here we characterize >100 autosomal loci with allele-specific epigenetic regulation of replication timing and gene expression, defining them as I/SCs. I/SCs are approximately 1 megabase in size and can contain both protein-coding and noncoding genes. In different single cell derived clones, these genes may be expressed from a single allele, the opposite allele, both alleles, or not expressed at all. This stochastic, yet mitotically stable, pattern indicates that the choice of which allele is expressed is independent of parent of origin and independent of the expression status of the other allele. Similarly, alleles within I/SCs show varying replication timing, either earlier or later, that is also independent of the other allele. Additionally, we identify syntenic loci in the mouse genome that display epigenetic regulation of allelic replication timing, highlighting the genomic organization and conservation of I/SC-associated regulation between human and mouse genomes. The allele-restricted regulation described here creates extensive cellular mosaicism through a stable epigenetic mechanism. This mosaicism impacts numerous dosage-sensitive genes associated with human diseases such as Alzheimer, Parkinson, epilepsy, deafness, and impaired intellectual development.

## Introduction

The relationship between monoallelic expression and asynchronous replication between alleles of autosomal genes, exemplified by genes subject to genomic imprinting or allelic exclusion, has been the focus of extensive research for over three decades ^2–4^. In recent work, we employed haplotype-phased RNA expression and replication timing assays on multiple single-cell-derived Lymphoblastoid Cell Line (LCL) clones from two unrelated individuals to identify loci exhibiting allele-restricted expression and replication timing ^1^. This clonal, allele-specific analysis revealed an unexpected genomic feature affecting megabase-sized loci across every autosome, characterized by allele-restricted gene expression (including random monoallelic expression) and variable replication timing (including asynchronous replication between alleles). The magnitude of allelic differences in expression and replication timing observed in some of the clones was comparable to the large differences detected between the active and inactive X chromosomes ^1^. We have named these autosomal loci Inactivation/Stability Centers (I/SCs) ^1,5–7^.

I/SCs can encompass both coding and noncoding genes that may be expressed from a single allele in some clones, expressed from the opposite allele in other clones, biallelically expressed in other clones, or silent on both alleles in yet other clones. This pattern of allele-restricted expression indicates that each allele independently adopts either an expressed or silent state. Importantly, because these expression states are mitotically stable, allele-autonomous, and independent of parental origin, we refer to the choice of the expressed allele as stochastic ^1^. To describe this phenomenon, we and others have used the term Allelic Expression Imbalance (AEI) to refer to the stochastic nature of autosomal random monoallelic expression ^1,8–11^. Additionally, the replication timing profiles on autosomes are highly variable and clone-specific, exhibiting both synchronous and asynchronous patterns in different clones. We refer to this as Variable Epigenetic Replication Timing (VERT) ^1^.

Random monoallelic expression in the olfactory, immune, and central nervous systems is well established and plays a critical role in generating cell-specific identities ^12–14^. However, increasing evidence indicates that autosomal genes outside these systems also exhibit allele-specific expression, suggesting that AEI plays a broader role across diverse cell types ^4,13^. The functional diversity of these genes implies that AEI contributes to a broad array of cellular processes and is not limited to specialized systems.

In the present study, we aimed to determine whether I/SCs could be detected in the genomes of clonal populations of human primary cells derived from a tissue distinct from those previously examined. To this end, we employed articular cartilage progenitor (ACP) cells as a second model system ^15^. ACP cells are non-transformed yet are particularly well-suited for clonal analysis due to several advantages: 1) they can be readily isolated from amputated tissue of patients with polydactyly, 2) they exhibit high proliferative capacity, and 3) they can be efficiently expanded from single cells ^16^. We identify an additional 163 VERT regions in the human genome, increasing the total number of potential I/SCs to 363, which collectively cover approximately 12% of the human genome. Of these, 112 loci display both AEI and VERT at the same genomic position and therefore represent high-confidence I/SCs.

Notably, we found that many previously reported “random monoallelic” genes, including those encoding olfactory and vomeronasal receptors^3,17^, antigen receptors^18^, and homophilic cell adhesion proteins^12^, are located within VERT regions. This confirms that these well-established monoallelically expressed genes reside within genomic loci that display the defining characteristics of I/SCs: allele-restricted expression and variable replication timing.

The relationship between genotype and phenotype is central to understanding human genetic disease. While most loss-of-function mutations are recessive, indicating that a single normal allele is sufficient for typical function, dominantly inherited disorders often result from gain-of-function mutations. However, a significant fraction of dominant diseases arises through haploinsufficiency (i.e. heterozygous for a loss of function mutation), where a single functional allele fails to maintain normal physiology. Although haploinsufficiency may affect up to 40% of human genes ^19–21^, the underlying molecular mechanisms remain poorly understood. Current models suggest that insufficient gene dosage may arise from a lack of redundancy, critical expression thresholds, stoichiometric imbalances, or toxicity associated with a lack of overexpression buffering ^22–25^. In contrast, insights from X-linked disease inheritance illustrate how cellular mosaicism and cellular selection can shape phenotypic outcomes: in XX individuals, differences in the survival or expansion of cells expressing either the wild-type or mutant alleles can influence whether a disease manifests as recessive or dominant ^21,26,27^.

The magnitude of AEI is often quantified using allelic expression ratios, which represent the proportion of transcripts derived from each allele. Importantly, recent findings have demonstrated that even modest allelic expression biases can have functional consequences, particularly in disease contexts. One recent study directly linked small but stable AEI to variable clinical outcomes among individuals carrying the same pathogenic mutation, underscoring AEI as a potential modulator of disease expressivity ^28^. These observations highlight the importance of considering AEI as a disease modifier that may shape phenotypic diversity and influence the penetrance of genetic disorders.

Here, we demonstrate that numerous genes implicated in autosomal human genetic diseases, such as Parkinson (DNAJC6, LRRK2 and SNCA), epilepsy (SCN1A, GABR1, and SAMD12), deafness (ATP11A, EPS8, and MYO6), and neurodevelopmental disorders (KCNA1, RETREG1, ROBO1, and TIAM1), are located within VERT regions and display AEI. Furthermore, mapping I/SCs across all autosomes reveals 87 known dominantly inherited single-gene disorders that map within VERT regions, where loss of function mutations in most of these genes result in haploinsufficiency ^29^. We propose that epigenetically regulated stochastic AEI can contribute to pathogenic phenotypes in heterozygous individuals by either reducing the number of functional cells below a critical disease threshold, akin to X linked dominant disorders in females, or by disrupting the normal mosaicism of gene expression within disease-relevant tissues.

## Results

Here, we employed articular cartilage progenitor (ACP) cells^15^ as a distinct primary cell type from those used in our previous studies, serving as a second model system to identify I/SCs. ACP cells were isolated from amputated digits of patients with polydactyly, and genomic DNA from their parents was sequenced to generate phased genomes for allele-specific analyses. We examined two independent sets of ACP clones derived from unrelated individuals, one female (ACP7) and one male (ACP6), using haplotype-phased Repli-seq and RNA-seq (Figure 1a and 1b).

**Figure 1:**
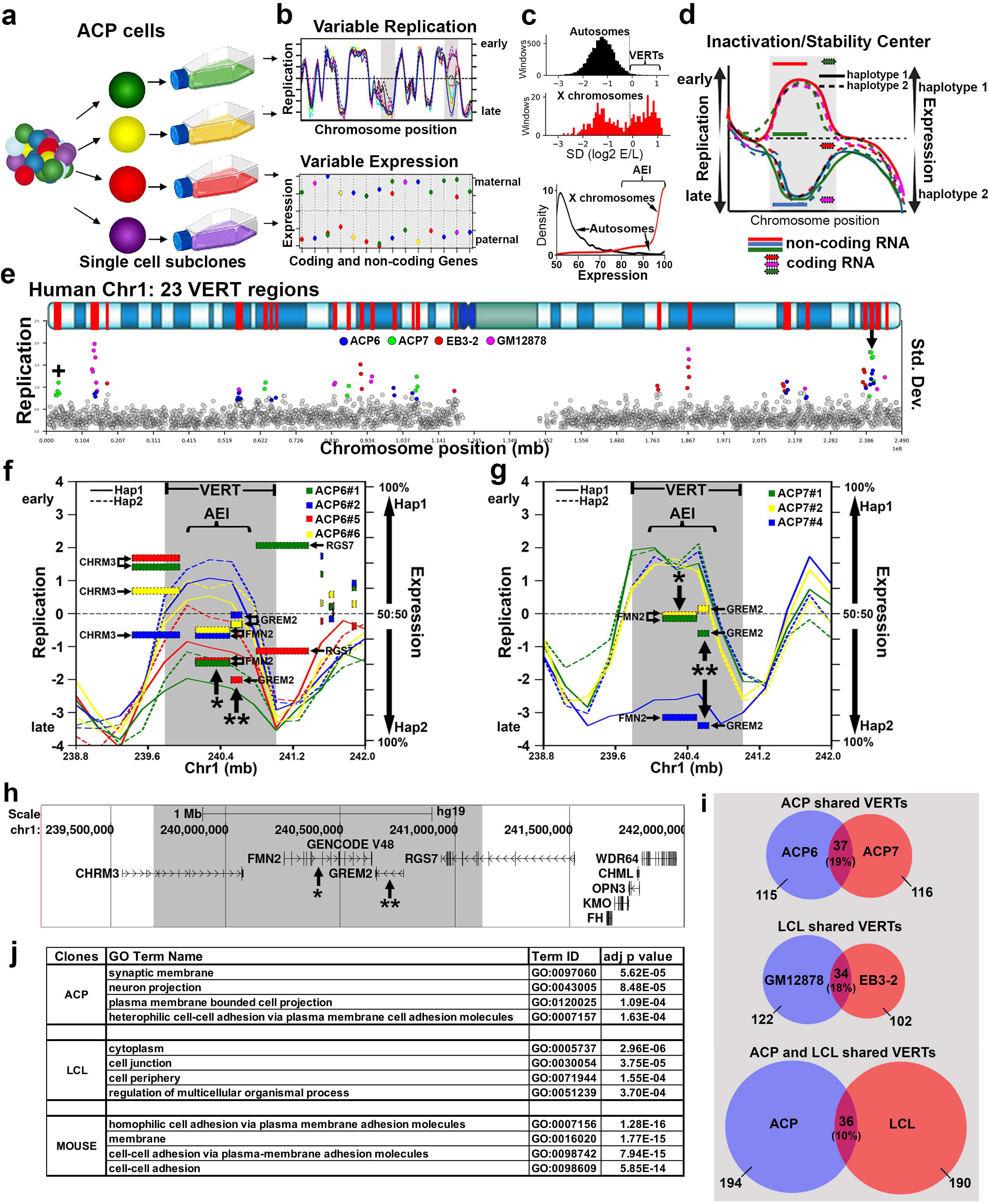
**Genome wide variable replication-timing and expression imbalance**. a) Single cell derived ACP clones (color coded) were isolated, expanded and processed for Repli-seq and RNA-seq. The maternal and paternal haplotypes were used to analyze allele-specific replication timing and expression. b) The top panel illustrates the expected replication timing profile in multiple clones across an autosomal region. The regions that display VERT are highlighted by shading. The bottom panel illustrates the expected expression variability in different clones from the same individual. c) We define VERT regions in Repli-seq data across each allele for each 250 kb genomic window. Windows with an SD value ≥ 2.5 standard deviations above the genome-wide median were classified as outliers (top panels). We define AEI as an allelic bias that is ≥80% allelic imbalance (AEI ≥ 0.80 or ≤ 0.20). The bottom panel shows both autosomal and X linked expression data. d) Illustration of an Inactivation/Stability Center showing the clonal and allele-restricted variability in replication timing and expression of both protein coding and noncoding genes. e) The standard deviation (Std. Dev.) in 250 kb windows (circles) across human chromosome 1. Outlier windows from all four sets of clones (ACP6, ACP7, EB3-2, and GM12878) are highlighted in different colors as shown. The top panel illustrates the G-banding (blue shading) pattern for human chromosome 1, with 23 VERT regions highlighted in red. The location of an imprinted region with asynchronous replication, containing the maternally expressed TP73 gene (geneimprint.com), is marked with an +. The arrow marks the VERT region shown in panels f-h below. f-h) I/SC located between 239 and 242 mb of chromosome 1 (see panel e). f) VERT (shaded region) and AEI of genes (rectangles) detected in the ACP6 clones, each clone was color coded as shown. For replication, the solid lines represent the paternal (Hap1) allele and the dotted line represents the maternal (Hap2) allele. g) VERT (shaded region) and AEI of genes (rectangles) detected in the ACP7 clones, each clone was color coded as shown. h) UCSC Genome Browser view of the genomic region in panels f and g. The shaded area highlights the VERT region. The asterisks mark the genes highlighted in panels f and g. i) Venn diagram illustrating the number of VERT regions that overlap between the different clonal populations. j) Gene Ontology (GO) enrichment analysis of the genes located within the VERT regions, the most significant hits for each set of clones, organized by cell type, are shown. GO analysis of the mouse genes located within the pre-B cell clone VERT regions are also shown.

To benchmark the magnitude of allele-specific epigenetic differences in replication timing and gene expression, we incorporated two internal controls. First, analysis of female ACP7 clones enabled direct comparison of autosomal allelic differences with those observed between the active and inactive X chromosomes within the same clones, thereby establishing statistical thresholds for VERT and AEI assignment (Figure 1c; Supplementary Tables 1–3). Second, we examined allelic replication timing at known imprinted loci and identified 12 imprinted regions displaying allelic replication imbalance that is comparable in magnitude to the VERT regions detected at non-imprinted loci. Imprinted regions with differences in allelic replication timing are highlighted in Figure 1e, Supplementary Figure 1, as well as in subsequent figures (Figures 2d, 3c, 6g, 7e, and 7g).

**Figure 2.**
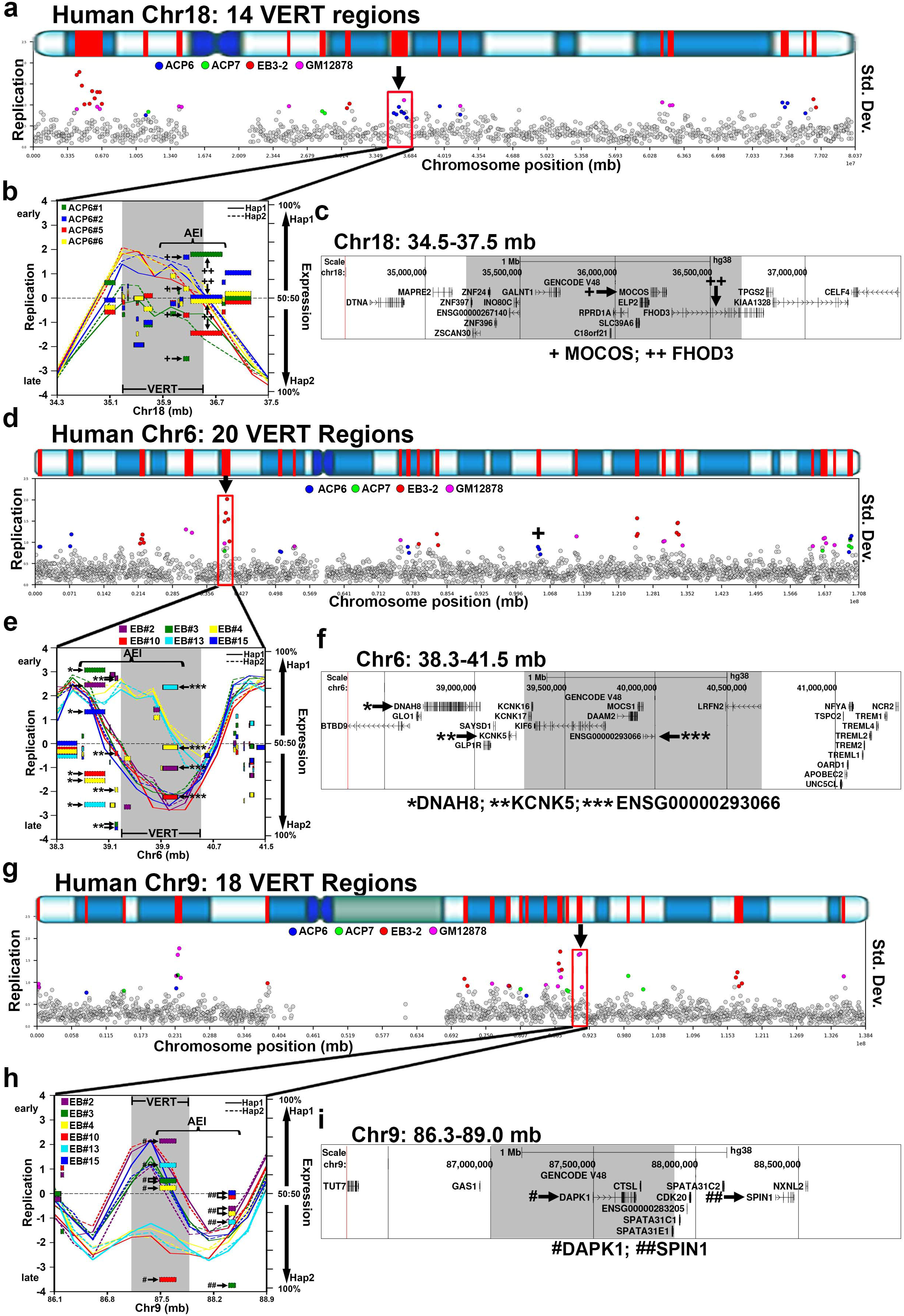
Examples of I/SCs. a, d and g) VERT regions on human chromosomes 18, 6 and 9, respectively. The top panels illustrate the G-banding (blue shading) pattern for each human chromosome, with VERT regions highlighted in red. The standard deviation (Std. Dev.) in 250 kb windows (circles) across each human chromosome is shown. Outlier windows from all four sets of clones (ACP6, ACP7, EB3-2, and GM12878) are highlighted in different colors as shown. The arrows and red boxes mark the VERT regions shown in panels b, e, and h. b, e and h) I/SCs highlighted in panels a, d, and g. For replication, the solid lines represent the paternal (Hap1) allele and the dotted lines represent the maternal (Hap2) allele. The regions with VERT are shaded and the AEI is shown for each clone in different colors. The location of an imprinted region with asynchronous replication, containing the paternally expressed LIN28B gene (geneimprint.com), is marked with an + in panel d. c, f and i) UCSC Genome Browser views of the genomic regions in panels b, e and h. The shaded areas highlight the VERT regions. Genes with AEI are marked with +, *, or ^#^. b and c) The + marks MOCOS, which shows AEI in ACP cells (Supplemental Table 2) and ++ marks FHOD3, which shows AEI in LCLs ^1^. e and f) The * marks from DNAH8, ** marks transcripts from KCNK5, and *** marks non-coding transcripts from ENSG00000293066 (also named: TL:6-39.9_1 ^1^). All three of these genes show AEI in LCL clones ^1^. h and i) The ^#^ marks transcripts from DAPK1, ^##^ marks transcripts from SPIN1. Both genes show AEI in LCL clones ^1^.

To identify genomic regions with epigenetic variation in replication timing between alleles, we calculated the standard deviation (SD) of Repli-seq data across each allele for each 250 kb genomic window. Windows with an SD value ≥ 2.5 standard deviations above the genome-wide median were classified as outliers. This analysis identified 163 additional VERT regions (Supplementary Table 1). Figure 1e illustrates this analysis for both ACP (ACP6 and ACP7) and LCL (EB3-2 and GM12878) clones on chromosome 1, where windows are indicated as gray circles and outliers indicated in different colors as shown.

Next, to detect coding genes with AEI in ACP cells, we employed RNA-seq on each clone and analyzed the number of reads at heterozygous SNPs. We defined AEI as differential expression between alleles using recently established criteria ^1,28^. Genes were considered AEI-positive if they exhibited >80% allelic imbalance (AEI ≥ 0.80 or ≤ 0.20) with FDR ≤ 0.05 in at least one clone. We selected the ≥80% allelic imbalance threshold as our criterion for significant AEI because a recent study demonstrated that allelic imbalance, as low as a 65%/35%, is enough to affect disease penetrance in humans ^28^. Additional analyses using more stringent thresholds (AEI ≥ 0.90 or ≥ 0.95) are provided in Supplementary Table 2. In addition, to exclude imprinted genes and effects due to genetic polymorphisms (i.e. expression Quantitative Trait Loci, eQTL^30^) we required that each AEI gene also be biallelically expressed (AEI < 0.70) in at least one clone from the same individual with ≥ 20 informative reads. Using these criteria, we identified 256 (2.4% of expressed genes) autosomal coding genes exhibiting AEI across the two ACP clone sets (Supplementary Table 2).

In addition, we also sought to identify AEI of extremely long noncoding RNAs (i.e. ASAR lincRNAs ^1,5,7,31^) expressed in the ACP clones. For this analysis we refer to regions of the genome with >50 kb of contiguous transcription of noncoding DNA as <u>T</u>ranscribed <u>L</u>oci (TL) ^1^. Using the same criteria used for the coding genes described above, we identified 23 TLs exhibiting AEI across the two ACP clone sets (Supplementary Table 3).

### Allelic Expression Imbalance and Variable Epigenetic Replication Timing define Inactivation/Stability Centers

By integrating the locations of genes exhibiting AEI with genomic regions displaying VERT, we identified overlapping loci that define I/SCs. A schematic representation of a hypothetical I/SC, showing a shaded VERT region in the replication profile and allele-specific expression of coding and noncoding genes, is provided in Figure 1d. Figures 1f and 1g illustrate a representative I/SC located on chromosome 1 at approximately 240 Mb (marked by an arrow in Figure 1e), where VERT was detected across both sets of ACP clones. Figure 1h shows the corresponding UCSC Genome Browser view of this locus, with genes displaying AEI indicated by asterisks (see Supplementary Table 2). Additional examples of I/SCs located on chromosomes 18, 6, and 9 are presented in Figure 2. In total, we identified 112 loci where both VERT and AEI were detected at the same genomic position, representing high-confidence I/SCs (Supplementary Table 4).

One of the objectives of this study was to assess whether different tissues utilize distinct sets of I/SCs, as might be expected given that each cell type expresses a unique repertoire of genes potentially subject to AEI. To explore this, we compared the genomic locations of VERT regions identified in the two ACP clonal sets and the two LCL clonal sets. As shown in Figure 1i, Venn diagrams reveal a modest degree of tissue-specific overlap: ACP6 clones share more VERT regions with ACP7 than with either LCL clone set, and the reciprocal relationship holds for the LCL clones. These findings suggest that while a subset of I/SCs are shared across tissues, a modest component of tissue-specific epigenetic regulation of allelic replication timing also exists.

At the molecular level, stochastic monoallelic expression contributes to cellular individuality and introduces functional variability among individual cells within complex biological systems ^12–14^. This mechanism is particularly well characterized in the olfactory, immune, and central nervous systems, where the expression of diverse receptor-type molecules through random monoallelic expression confers cell-specific functional identities. To further investigate the functional characteristics of protein-coding genes located within VERT regions, we annotated the genes residing within the VERT regions identified across all four clonal datasets (ACP clones: ACP6 and ACP7; and LCL clones: EB3-2, and GM12878; see Supplementary Table 5). To assess whether these genes are functionally related, we performed a Gene Ontology (GO) enrichment analysis. This analysis revealed a significant enrichment for genes involved in plasma membrane localization, cell-cell adhesion and neurodevelopmental processes, the most significant hits for each set of clones are presented in Figure 1j (see full details in Supplementary Table 6). This analysis suggests that I/SC-associated epigenetic regulation may play a broader role in the establishment of cellular identity and tissue organization particularly in the nervous system.

### Gene clusters with AEI map to VERT regions

In the olfactory, immune, and central nervous systems, random monoallelic expression of gene family members, often organized in genomic clusters, plays a key role in establishing cell-specific identities ^12–14^. Consistent with these observations, we found that VERT regions frequently coincide with gene clusters that contain known AEI genes (Table 1). For example, Figure 3a and 3b highlight a cluster of 11 olfactory receptor (OR) genes located on human chromosome 19 at approximately 15 Mb. This region also harbors a cluster of Cytochrome P450 (CYP4F family, n = 6) genes. The interspersed genomic arrangement of the CYP4F genes among the OR genes, indicates that both gene families reside within the same I/SC and therefore both may be subject to AEI.

**Figure 3.**
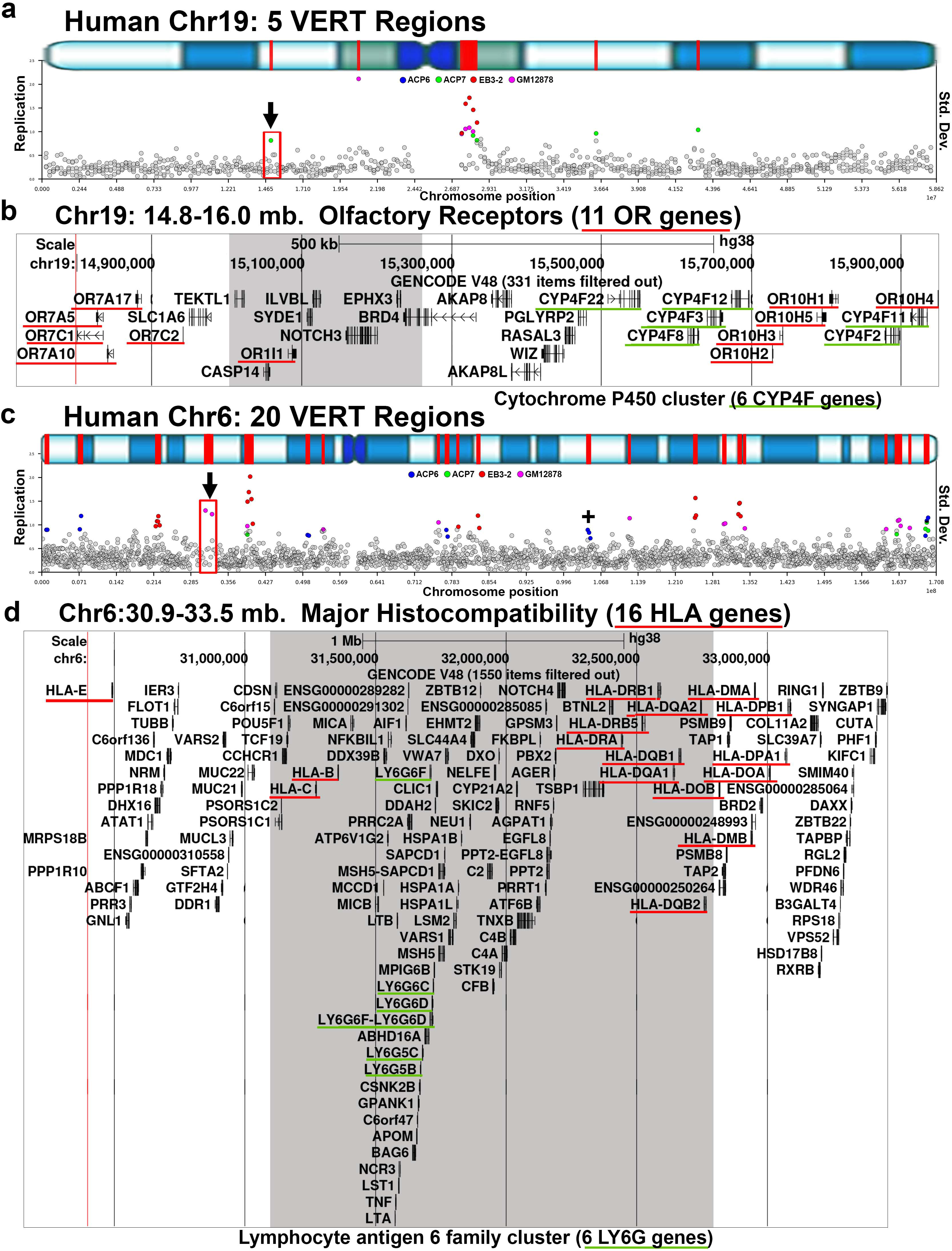
Human gene clusters with AEI map to VERT regions. a and c) VERT regions on human chromosome 19 and 6, respectively. The top panels illustrate the G-banding (blue shading) pattern for each human chromosome, with VERT regions highlighted in red. The standard deviation (Std. Dev.) in 250 kb windows (circles) across each human chromosome is shown. Outlier windows from all four sets of clones (ACP6, ACP7, EB3-2, and GM12878) are highlighted in different colors as shown. The red boxes and arrows identify the I/SCs in panel b and d. The location of an imprinted region with asynchronous replication, containing the paternally expressed LIN28B gene (geneimprint.com), is marked with an + in panel c. b) UCSC genome browser view illustrating the genomic location highlighted in panel a. The shaded area highlights the VERT region. The red underlines mark 11 olfactory receptor genes, and the green underlines mark 6 cytochrome P450 family (CYP4F) genes. d) UCSC Genome Browser view illustrating the genomic location highlighted in panel c. The shaded area highlights the VERT region. The red underlines mark 16 HLA genes, and the green underlines mark 6 Lymphocyte antigen 6 family (LY6G) genes.

**Table 1.** VERT regions at gene clusters in the human and mouse genomes.

| Human VERT Region |  |  | Gene Cluster |  |  |  |
| --- | --- | --- | --- | --- | --- | --- |
| Chr | Start | Stop | Genes | # of Genes | Start | stop |
| 1 | 13,250,000 | 15,000,000 | PRAMEF | 23 | 12,768,939 | 13,422,488 |
| 2 | 101,633,538 | 102,383,540 | IL1R | 5 | 101,987,899 | 102,401,443 |
| 2 | 165,250,000 | 166,000,000 | SCN | 5 | 165,081,204 | 166,499,088 |
| 2 | 176,250,000 | 176,500,000 | HOXD | 9 | 176,091,062 | 176,191,493 |
| 4 | 46,750,000 | 47,500,000 | GABR | 4 | 46,035,583 | 47,431,569 |
| 4 | 68,384,282 | 68,634,282 | UGT | 9 | 68,525,283 | 69,660,413 |
| 5 | 651,700 | 750,000 | SLC | 5 | 468,888 | 1,448,927 |
| 5 | 148,000,000 | 148,250,000 | SPINK | 7 | 147,822,992 | 148,343,066 |
| 5 | 162,250,000 | 162,750,000 | GABR | 4 | 161,282,809 | 162,159,566 |
| 6 | 31,250,000 | 32,750,000 | HLA | 15 | 30,489,509 | 33,093,613 |
|  |  |  | LY6 | 6 | 31,669,259 | 31,722,476 |
| 8 | 11,250,000 | 12,392,491 | DEFB | 6 | 11,970,747 | 12,320,078 |
| 8 | 80,587,765 | 83,000,000 | FABP | 4 | 81,277,942 | 81,536,065 |
| 8 | 142,500,000 | 142,750,000 | LY6 | 8 | 142,695,707 | 143,162,219 |
| 11 | 104,750,000 | 105,000,000 | CASP | 5 | 104,883,181 | 105,141,441 |
| 15 | 23,357,400 | 23,500,000 | GOLGA | 5 | 22,451,995 | 23,451,545 |
| 15 | 28,750,000 | 29,000,000 | GOLGA | 6 | 28,344,604 | 28,855,059 |
| 17 | 9,750,000 | 10,500,000 | MYH | 6 | 10,297,552 | 10,658,865 |
| 19 | 15,000,000 | 15,250,000 | OR | 11 | 14,792,476 | 15,952,060 |
|  |  |  | CYP | 6 | 15,507,105 | 15,936,041 |
| 19 | 20,750,000 | 21,000,000 | ZNF | 37 | 19,667,773 | 24,119,420 |
| 19 | 36,313,700 | 36,500,000 | ZNF | 29 | 36,171,632 | 37,791,237 |
| 19 | 43,000,000 | 43,250,000 | PSG | 10 | 42,713,521 | 43,272,861 |
| 20 | 24,250,000 | 24,750,000 | CST | 11 | 23,429,902 | 24,957,423 |
| 21 | 30,627,681 | 30,877,681 | KRTAP | 33 | 30,276,201 | 31,041,124 |

| Mouse VERT Region |  |  | Gene Cluster |  |  |  |
| --- | --- | --- | --- | --- | --- | --- |
| Chr | Start | Stop | Genes | # of Genes | Start | stop |
| 2 | 36,322,857 | 36,622,857 | Or | 28 | 36,225,624 | 37,193,679 |
| 2 | 64,622,866 | 66,522,868 | Scn | 5 | 65,287,042 | 66,637,709 |
| 2 | 88,950,000 | 89,950,000 | Or | 239 | 85,160,859 | 90,147,389 |
| 4 | 59,850,000 | 59,950,000 | Mup | 22 | 59,950,000 | 61,824,646 |
| 5 | 91,150,000 | 91,350,000 | Cxcl | 5 | 91,186,938 | 91,335,955 |
| 5 | 95,950,000 | 96,250,000 | Pramel | 23 | 93,891,822 | 96,243,866 |
| 6 | 40,650,000 | 41,150,000 | Tas2r | 8 | 40,386,430 | 43,196,843 |
|  |  |  | Or | 24 | 40,386,430 | 43,196,843 |
| 6 | 66,650,000 | 66,850,000 | Vmn1 | 8 | 66,498,911 | 66,758,688 |
| 7 | 113,809,945 | 113,909,945 | Or | 24 | 113,790,926 | 114,306,041 |
| 11 | 73,450,000 | 74,050,000 | Or | 31 | 73,160,403 | 74,180,092 |
| 18 | 37,350,000 | 37,750,000 | Pcdh | 23 | 37,298,052 | 37,684,088 |

A second example of a gene cluster is shown in Figure 3c and 3d, where the <u>m</u>ajor <u>h</u>istocompatibility <u>c</u>omplex (MHC), comprising 16 HLA genes (which are known to be subject to AEI ^28,32^), lies within a VERT region. Notably, this locus also contains a cluster of Lymphocyte Antigen 6 genes (LY6G family, n = 6), which are interspersed among the HLA genes. This genome organization indicates that the LY6G genes, like their neighboring HLA genes, are within the same I/SC and suggests that they too may be subject to stochastic allelic expression imbalance.

### Mapping mouse VERTs identifies known AEI gene clusters and syntenic I/SCs

Much of our current understanding of stochastic autosomal allelic expression is derived from studies conducted in the mouse ^4,9–11,17,18,33–40^. To assess whether VERT regions could be identified at expected loci in the mouse genome, specifically at sites of known AEI, we analyzed previously published Repli-seq data from mouse pre-B cell clones reported by Blumenfeld et al ^41^. This analysis revealed 117 loci in the mouse genome that display VERT (Supplemental Table 7).

Consistent with our findings in the human genome, the VERT regions detected in the mouse genome frequently overlap with genes that are known to display AEI, including 5 olfactory receptor gene clusters located on four different autosomes (Table 1). Figure 4 presents three examples: two olfactory receptor (Or) gene clusters on mouse chromosome 2 (Figure 4a–4c) and the protocadherin (Pcdh) gene cluster, which are known to generate cellular individuality in the central nervous system through stochastic expression of different Pcdh family members ^12,42^, located on mouse chromosome 18 (Figure 4d–4e). In addition to these gene clusters with VERT, we also detected immunoglobulin and T cell receptor gene loci with VERT. As detailed in Supplemental Figure 2, we detected VERT at the T cell receptor alpha (Tra) locus on mouse chromosome 14 at approximately 54 Mb. One striking feature of this locus is that it also contains an olfactory receptor cluster, containing 37 Or genes, and a vomeronasal receptor gene (Vmn2r88). All of these multigene families are well documented sites of AEI ^3,12,17,42,43^, highlighting the genomic organization and conservation of I/SC-associated regulation at known AEI genes in the mouse. Gene Ontology analysis of the mouse genes located within VERT regions showed significant enrichment for biological processes related to the plasma membrane and cell-cell adhesion (Figure 1j), which is consistent with the human VERT gene dataset and reinforces the conservation of this regulatory mechanism across species.

**Figure 4.**
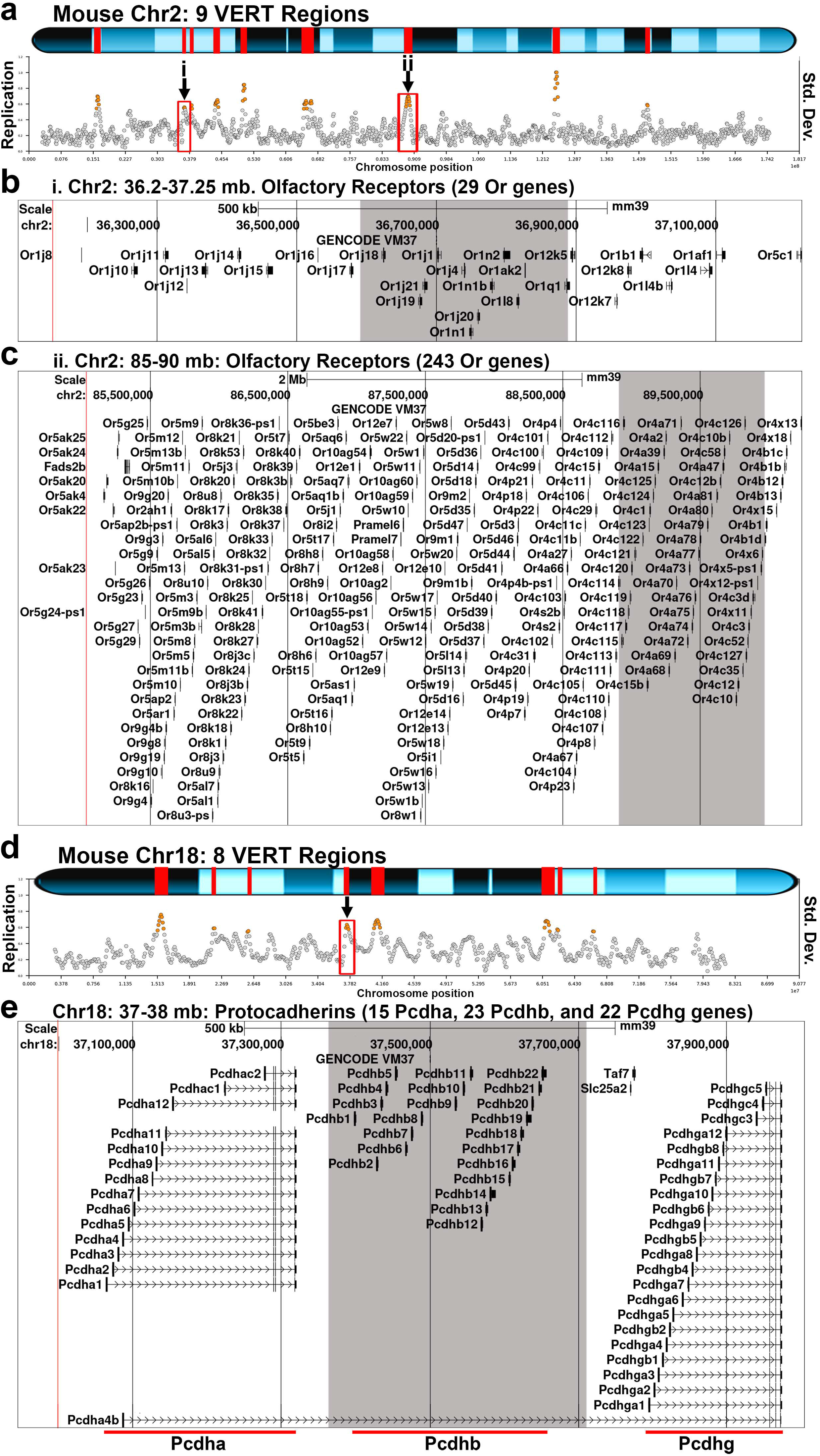
Mouse gene clusters with AEI map to VERT regions. a and d) VERT regions on mouse chromosome 2 and 18, respectively. The standard deviation (Std. Dev.) in 50 kb windows (circles) across each mouse chromosome are shown. Outlier windows from pre-B cell clones are highlighted in orange. The red boxes and arrows identify the VERT regions in panels b (i) and c (ii). b and c) UCSC genome browser views illustrating the genomic locations represented by i and ii in panel a. b) Genome Browser view of the VERT region showing the location of 29 olfactory receptor genes (Or genes). The shaded area represents the VERT region. c) Genome Browser view of the VERT region in ii above, showing the location of 242 olfactory receptor genes (Or genes). The shaded area represents the VERT region. e) Genome Browser view of the VERT region in panel d, showing the location of clustered Protocaderhin genes (15 Pcdha, 23 Pcdhb, and 22 Pcdhg genes). The shaded area represents the VERT region.

Previous studies analyzed a relatively small number of mouse AEI genes to model their human counterparts with respect to autosomal-dominant disorders ^38,44^. To further investigate the conservation of I/SC regulation between the human and mouse genomes, we identified VERT regions that map to syntenic loci across the two species. This analysis revealed 21 loci where VERT is detected in both genomes, as detailed in Table 2. Figure 5 presents three representative examples of syntenic VERT regions: two loci on the short arm of human chromosome 5 with corresponding regions on mouse chromosome 15 (Figs. 5a–5g), and a third locus on human chromosome 16 with its syntenic region on mouse chromosome 16 (Figs. 5h–5k). Additional examples of syntenic loci showing AEI and/or VERT, including mouse App (see Gimelbrant et al. ^45^ for AEI of human APP), Mocos and Fhod3 (see Figure 2a for VERT and AEI of human MOCOS and FHOD3), and a cluster of 5 sodium channel genes, Scn1a, Scn2a, Scn3a, Scn7a and Scn9a (see Figure 6a-c below for VERT and AEI of the syntenic human SCN gene cluster) are shown in Supplemental Figure 3.

**Figure 5.**
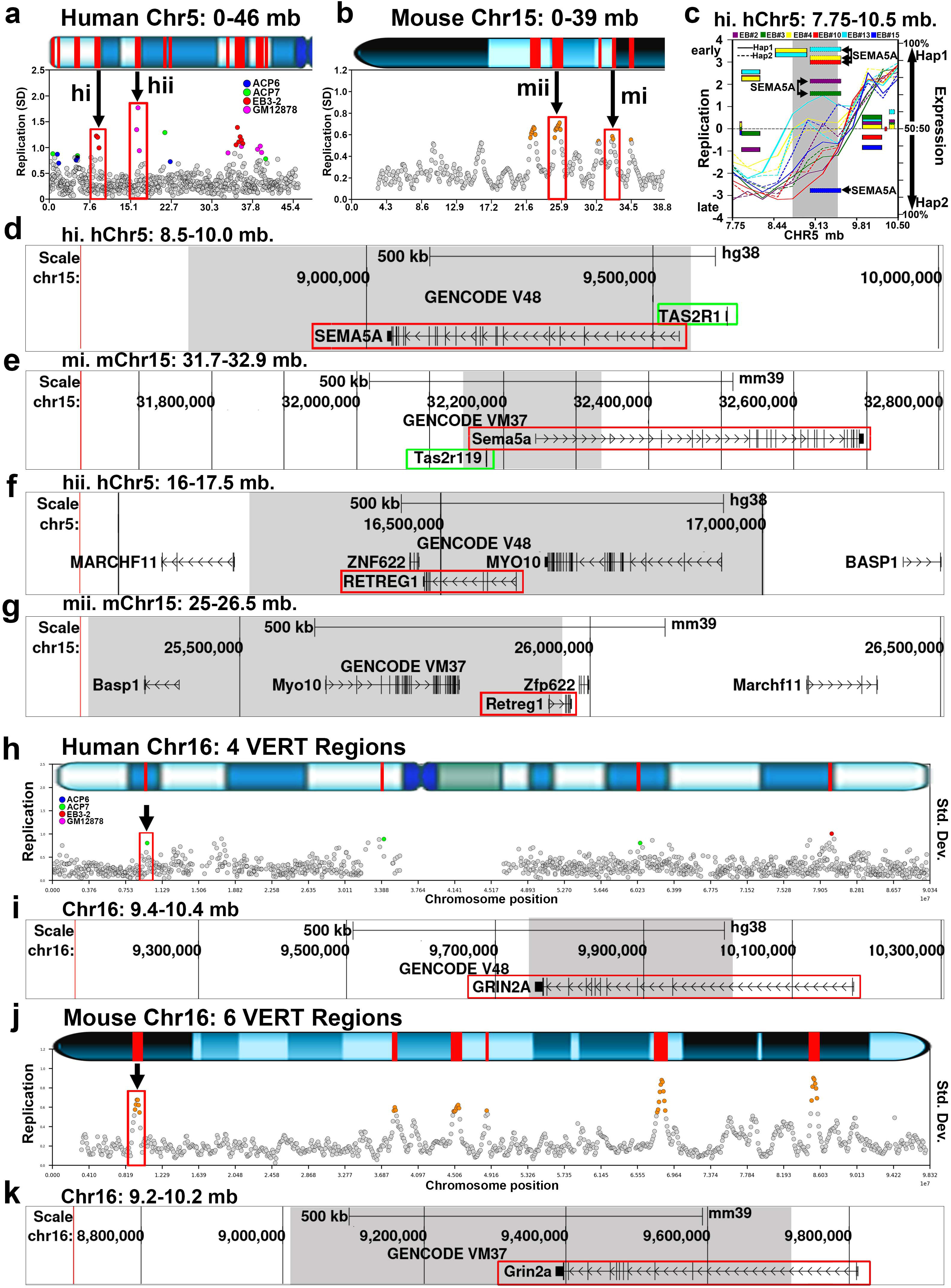
Human and Mouse syntenic regions display VERT. a and b) The short arm of human chromosome 5, with 11 VERT regions (panel a) and the centromeric region of mouse chromosome 15, with 4 VERT regions (panel b) are shown as orange circles. a) Two human VERT regions are highlighted, hi and hii, and are syntenic with locations on mouse chromosome 15, mi and mii, that also show VERT (panel b). Note that the gene order in both syntenic regions are inverted with respect to the telomeres when comparing the human and mouse loci. c) Illustration of the I/SC on human chromosome 5 between 7.75 and 10.5 mb (hi). The shaded area marks the VERT region, and AEI of the coding gene SEMA5A in different clones is highlighted. For replication, the paternal (Hap1) and maternal (Hap2) alleles are indicated. d) UCSC Genome Browser view of the human VERT region (shaded) in hi, showing the location of SEMA5A and a taste receptor gene (TAS2R1). e) UCSC Genome Browser view of the mouse VERT region (shaded) in mi, showing the location of Sema5a and a taste receptor gene (Tas2r119). f) UCSC Genome Browser view of the human VERT region (shaded) in hii above, highlighting the location of the AEI gene RETREG1 ^45^. g) UCSC Genome Browser view of the mouse VERT region (shaded) in mii, showing the synteny between human and mouse for the protein coding genes in panel f, including Retreg1. h) VERT regions on human chromosome 16. The standard deviation (Std. Dev.) in 250 kb windows (circles) across human chromosome 16 is shown. Outlier windows are highlighted in different colors as shown. The red box and arrow mark the genomic location in panel i. i) UCSC Genome Browser view illustrating the genomic location represented in panel h, showing the location of GRIN2A (red box). j) VERT regions on mouse chromosome 16. The standard deviation in 50 kb windows (circles) is shown. Outlier windows from pre-B cell clones are highlighted in orange. k) UCSC Genome Browser view illustrating the genomic location of the VERT region (shaded area) represented in panel j above, with the location of Grin2a highlighted with a red box.

**Figure 6.**
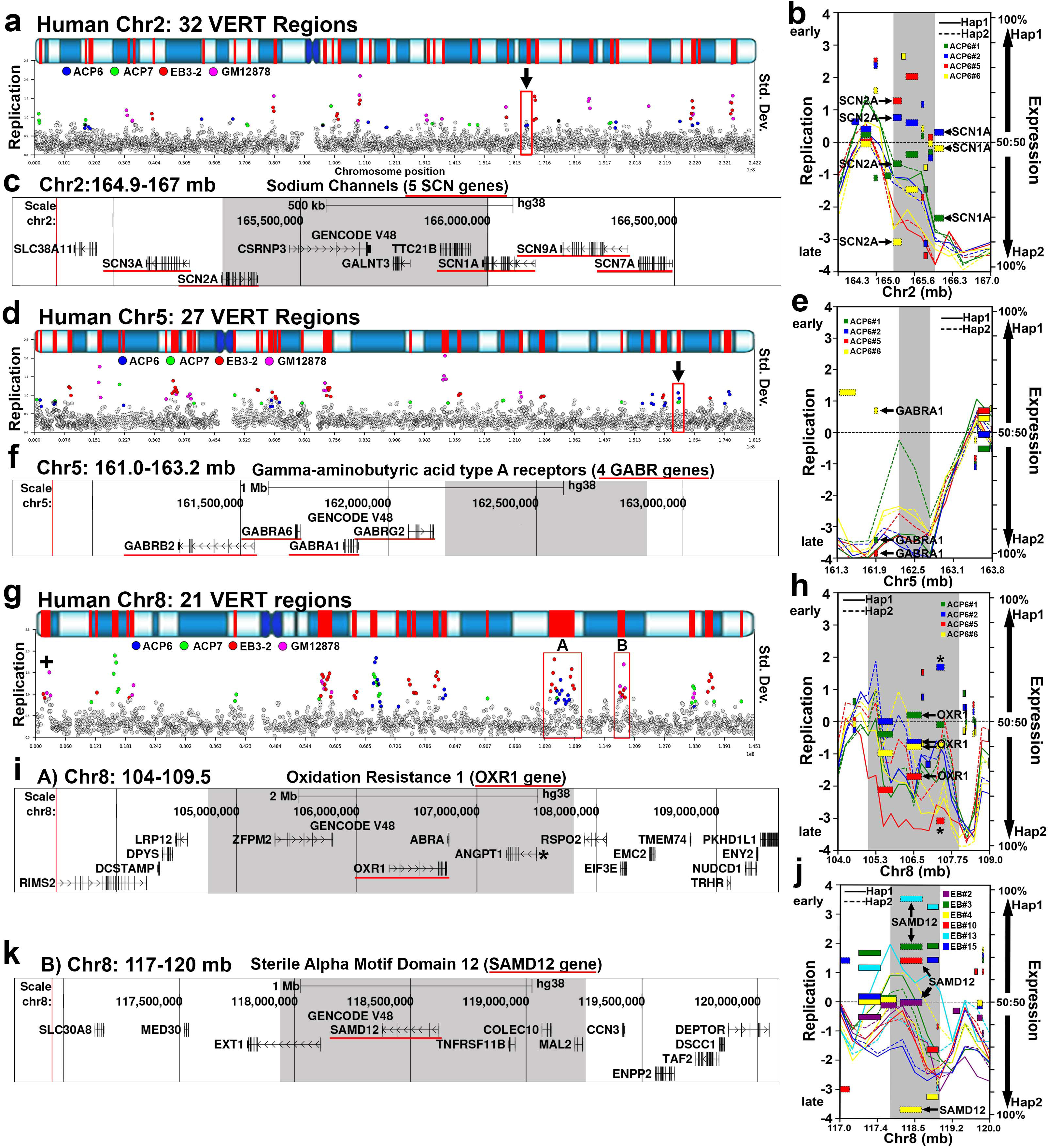
Epilepsy genes map to I/SCs. a, d, and g) VERT regions on human chromosomes 2, 5, and 8, respectively. The top panels illustrate the G-banding (blue shading) pattern for each human chromosome, with VERT regions highlighted in red. The standard deviation (Std. Dev.) in 250 kb windows (circles) across each human chromosome is shown. Outlier windows from all four sets of clones (ACP6, ACP7, EB3-2, and GM12878) are highlighted in different colors as shown. The location of an imprinted region with asynchronous replication, containing the paternally expressed DLGAP2 gene (geneimprint.com), is marked with an + in panel g. The arrows and red boxes mark the VERT regions shown in panels b, c, e, f, h, i, j and k. b, e, h, j) Illustrations of I/SCs on chromosomes 2, 5, and 8, respectively. The shaded areas mark the VERT regions. Each clone was color coded to illustrate the VERT and AEI in different clones. The paternal (Hap1) and maternal (Hap2) alleles are indicated for replication and expression. c) UCSC Genome Browser view illustrating the genomic location of the VERT region (shaded area) represented in panels a and b, with the location of 5 sodium channel genes (SCN3A, SCN2A, SCN1A, SCN9A, and SCN7A) highlighted with red underlines. e) Illustration of an I/SC on chromosome 5. The shaded area marks the VERT region. AEI of GABRA1 is shown. f) UCSC Genome Browser view illustrating the genomic location of the VERT region (shaded area) represented in panels d and e above, with the location of 4 GABA receptor genes (GABRB2, GABRBA6, GABRA1, and GABRG2) highlighted with red underlines. h) Illustration of an I/SC (A and red box) on chromosome 8. Each ACP6 clone was color coded to illustrate the VERT and AEI in different clones. The paternal (Hap1) and maternal (Hap2) alleles are indicated for replication and expression. The shaded area marks the VERT region. The location and expression of OXR1 is indicated. i) UCSC Genome Browser view illustrating the genomic location of the VERT region (shaded area) represented in panels g (A) and h above, with the location of the OXR1 gene highlighted with a red underline. We note that ANGPT1 (marked with *), located within this I/SC displays AEI (see Supplementary Table 2). j) Illustration of an I/SC (B and red box) on chromosome 8. AEI of SAMD12 is indicated. k) UCSC Genome Browser view illustrating the genomic location of the VERT region (shaded area) represented in panels g (B) and j above, with the location of the SAMD12 highlighted with a red underline.

**Table 2.** VERT regions showing synteny between human and mouse genomes.

| Mouse VERT |  |  | Synteny (%) | Human Syntenic Region |  |  | Human VERT |  |  |
| --- | --- | --- | --- | --- | --- | --- | --- | --- | --- |
| Chr | Start | stop |  | Chr | start | Stop | Chr | start | Stop |
| 1 | 73,042,585 | 73,542,585 | 54.0 | 2 | 216,836,864 | 217,400,721 | 2 | 217,000,000 | 217,250,000 |
| 1 | 88,681,147 | 89,081,147 | 33.0 | 2 | 234,565,154 | 235,053,857 | 2 | 234,000,000 | 235,000,000 |
| 1 | 189558318 | 189,758,318 | 49.8 | 1 | 214,144,079 | 214,410,956 | 1 | 214,326,657 | 216,000,000 |
| 2 | 64,842,287 | 66,742,287 | 46.5 | 2 | 164,610,983 | 166,685,353 | 2 | 165,250,000 | 166,000,000 |
| 3 | 18,204,164 | 18,404,164 | 33.7 | 8 | 64,684,742 | 64,876,026 | 8 | 64,337,443 | 65,087,765 |
| 3 | 57,253,499 | 58,553,499 | 32.6 | 3 | 149,407,811 | 150,704,641 | 3 | 149,282,213 | 150,032,213 |
| 3 | 78,253,385 | 78,653,385 | 26.1 | 4 | 159,655,594 | 159,952,325 | 4 | 159,328,848 | 159,828,848 |
| 3 | 82,453,385 | 82,853,385 | 32.6 | 4 | 154,688,934 | 155,210,940 | 4 | 154,500,000 | 154,750,000 |
| 4 | 105034592 | 105,234,592 | 62.8 | 1 | 56,331,724 | 56,556,339 | 1 | 55,500,000 | 56,750,000 |
| 4 | 142286667 | 142,686,667 | 40.6 | 1 | 14,022,915 | 14,523,879 | 1 | 13,250,000 | 15,000,000 |
| 8 | 10,500,000 | 11,200,000 | 43.2 | 13 | 108,978,718 | 110,065,357 | 13 | 109,250,000 | 109,500,000 |
| 8 | 24,029,544 | 24,929,544 | 37.3 | 8 | 40,152,844 | 41,090,451 | 8 | 40,250,000 | 40,750,000 |
| 8 | 39,817,687 | 40,717,687 | 28.6 | 8 | 15,936,545 | 16,964,053 | 8 | 15,750,000 | 17,000,000 |
| 9 | 116969950 | 117,469,950 | 42.3 | 3 | 28,837,647 | 29,402,683 | 3 | 29,000,000 | 29,500,000 |
| 11 | 46,927,325 | 48,127,325 | 35.2 | 5 | 155,465,678 | 156,698,430 | 5 | 155,500,000 | 156,250,000 |
| 12 | 33,715,134 | 33,815,134 | 36.1 | 7 | 19,330,767 | 19,416,599 | 7 | 19,250,000 | 19,500,000 |
| 12 | 105278049 | 105,578,049 | 41.6 | 14 | 95,853,630 | 96,269,387 | 14 | 96,000,000 | 96,250,000 |
| 13 | 11,095,964 | 11,672,619 | 29.9 | 1 | 237,711,340 | 238,470,922 | 1 | 237,604,200 | 238,750,000 |
| 15 | 25,120,331 | 25,920,331 | 42.9 | 5 | 16,534,412 | 17,724,462 | 5 | 16,250,000 | 17,000,000 |
| 15 | 32,220,391 | 32,420,391 | 60.7 | 5 | 9,377,059 | 9,575,503 | 5 | 8,749,888 | 9,499,888 |
| 16 | 9,067,771 | 9,667,771 | 41.6 | 16 | 9,476,924 | 10,028,307 | 16 | 9,750,000 | 10,000,000 |

We point out that many of the mouse genes that are located within VERT regions are associated with various human genetic diseases (Supplemental Table 8). One of the loci highlighted in Figure 5a-5d labeled “hi”, exhibits both VERT and AEI (Figure 5c and ^1^), affecting the protein-coding gene SEMA5A, a gene implicated in autism and neurodevelopmental disorders ^46–48^. Notably, this locus also contains a syntenic pair of taste receptor genes: TAS2R1 in humans and Tas2r119 in mice. These genes are part of the taste receptor family, which are known to harbor chromatin marks characteristic of AEI-regulated loci ^49^. Consistent with this observation, we also detected VERT at a taste receptor gene cluster (Tas2r) on mouse chromosome 6 (Table 1).

A second example of synteny is shown in Figure 5f-5g, where human RETREG1 and mouse Retreg1 are located within VERT regions. Human RETREG1 is a known AEI gene ^45^, and mutations in RETREG1 cause sensory and autonomic neuropathy ^29^. A third example is shown in Figure 5h-5k, where both human GRIN2A and mouse Grin2a are located within VERT regions, and loss of function mutations in GRIN2A in humans and Grin2a in mice, are known to cause autosomal dominant (i.e. haploinsufficiency) epilepsy ^29,50,51^. These findings indicate that I/SC-associated epigenetic regulation is conserved across species, affects functionally related genes and gene families, and may influence both human and mouse genetic inheritance patterns (see below).

### Neurodevelopmental disease genes map to I/SCs

To further investigate the relationship between I/SC regulation and human disease genes, we annotated the human protein-coding genes located within VERT regions (see Supplemental Table 5). This analysis identified 979 genes overlapping VERT regions, of which 272 are associated with single-gene disorders according to the OMIM database ^29^. Among these genes, 155 are linked to autosomal recessive diseases, 87 to autosomal dominant diseases, and 30 exhibit both dominant and recessive inheritance patterns (Supplemental Table 9). Consistent with our GO analysis, which revealed an enrichment for neurodevelopmental processes (see Figure 1j), we found that various neurodevelopmental disease genes were prominently represented among the disease-associated genes (Table 3). Moreover, it was not unexpected to observe gene clusters associated with neurodevelopmental disorders residing within VERT regions. For example, Figure 6a–6c depicts the location, I/SC profile, and genomic organization of a cluster of sodium channel genes (SCN family, n = 5). Mutations in SCN1A, SCN2A, SCN3A, and SCN9A genes are known to cause autosomal dominant developmental and epileptic encephalopathies, with haploinsufficiency as a primary mechanism (Table 3 and ^29,52,53^). We also note that the syntenic locus in the mouse displays VERT, with the 5 syntenic mouse Scn genes residing within a VERT region on mouse chromosome 2 (Supplementary Figure 3e and 3f).

**Table 3.**
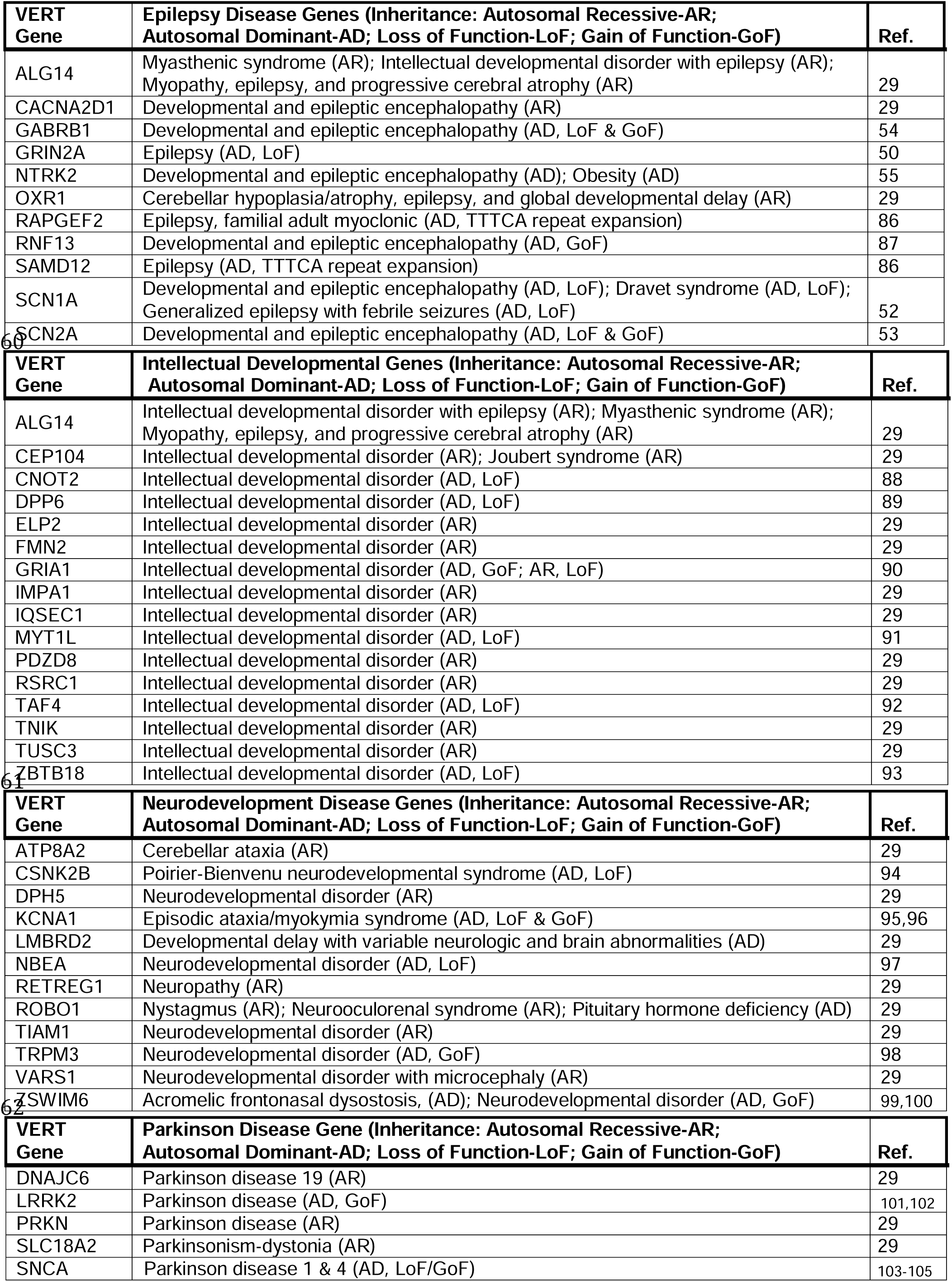
Neurodevelopmental genes within VERT regions.

A second example of a cluster of epilepsy disease genes is shown in Figure 6d–6e, where four members of the gamma-aminobutyric acid type A receptor gene family (GABR family, n = 4) reside adjacent to a VERT region; notably, GABRA1 displays AEI (Figure 6e, and Supplementary Table 2), and mutations in GABRA1, GABRB2, and GABRG2 cause autosomal dominant developmental and epileptic encephalopathies ^29,54,55^. We also note that a second GABR gene cluster (n = 4), where GABRA2 and GABRB1 are also associated with autosomal dominant developmental and epileptic encephalopathies, is adjacent to a VERT region on chromosome 4 (see Table 1). Two additional examples are shown in Figure 6g–6j, where OXR1 and SAMD12 are located in different VERT regions on chromosome 8, and mutations in each result in autosomal dominant epilepsy ^29^.

Another prominent group of human disease genes located within VERT regions are those associated with Parkinson disease (Table 3). As shown in Figure 7, four well-characterized Parkinson disease genes, SNCA, LRRK2, DNAJC6, and PARKN, are located within or adjacent to VERT regions. Importantly, both SNCA and LRRK2 were among the first genes identified as displaying AEI in the study by Gimelbrant et al. in 2007 ^45^. In our own dataset, we found that DNAJC6 exhibits AEI in LCL clones ^1^. We also highlight that JAK1, a gene involved in immune signaling, lies within the same I/SC as DNAJC6 (Figure 7f). JAK1 is a known AEI gene ^56,57^, and mutations in JAK1 cause autosomal dominant autoinflammation, immune dysregulation, and eosinophilia (AIIDE). Importantly, a recent study indicated that AEI of wild-type and mutant JAK1 alleles correlates with variable disease penetrance ^28^.

**Figure 7.**
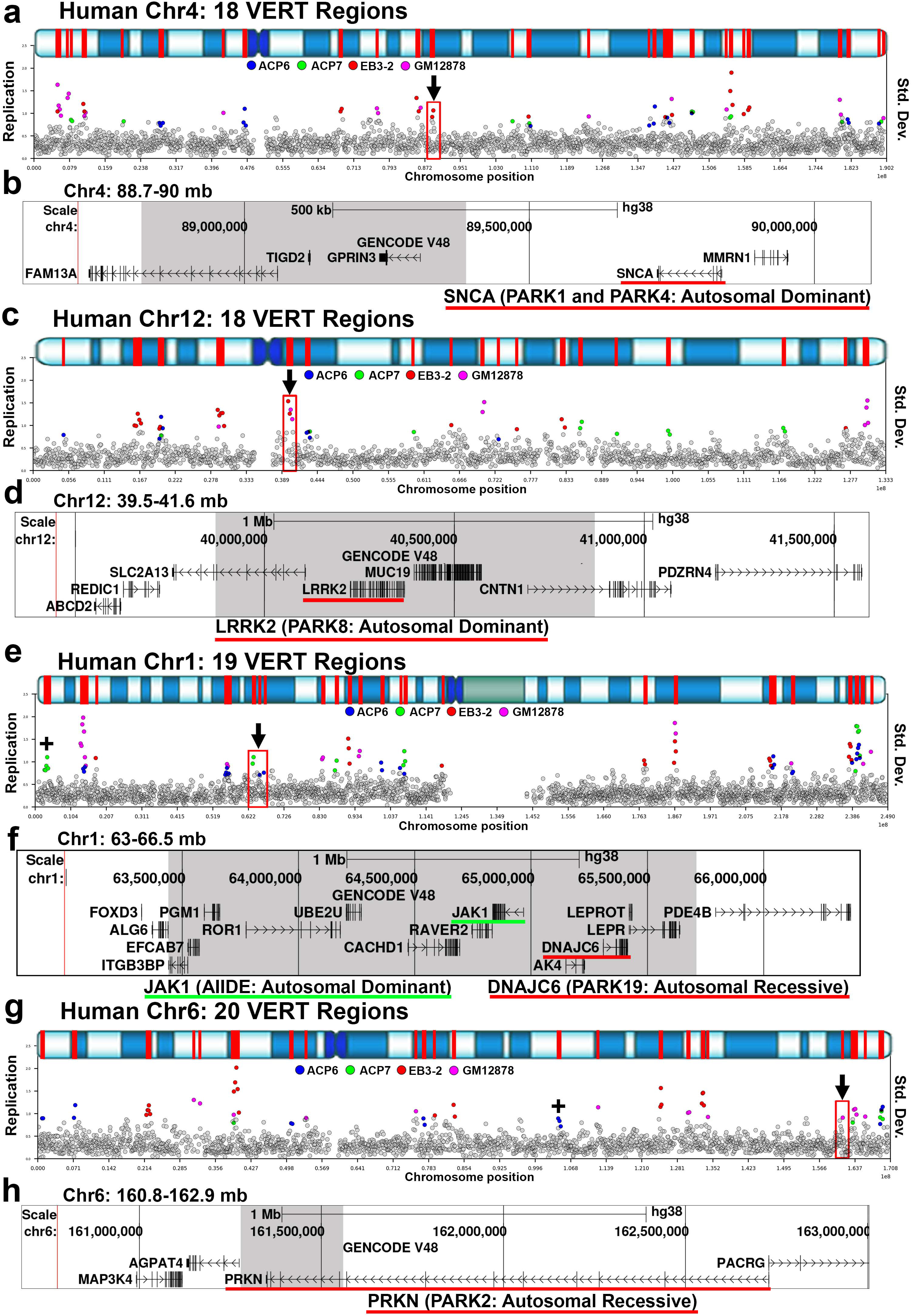
Parkinson disease genes map to I/SCs. a, c, e, and g) VERT regions on human chromosomes 4, 12, 1, and 6, respectively. The top panels illustrate the G-banding (blue shading) pattern for each human chromosome, with VERT regions highlighted in red. The standard deviation (Std. Dev.) in 250 kb windows (circles) across each human chromosome is shown. Outlier windows from all four sets of clones (ACP6, ACP7, EB3-2, and GM12878) are highlighted in different colors as shown. The location of an imprinted region with asynchronous replication, containing the paternally expressed LIN28B gene (geneimprint.com), is marked with an + in panel g. The location of an imprinted region with asynchronous replication, containing the maternally expressed TP73 gene (geneimprint.com), is marked with an + in panel e. The arrows and red boxes mark the VERT regions shown in panels b, d, f, and h, respectively. b) UCSC Genome Browser view illustrating the genomic location of the VERT region (shaded area) represented in panel a, with the location of the SNCA gene highlighted with a red underline. d) UCSC Genome Browser view illustrating the genomic location of the VERT region (shaded area) highlighted in panel c, with the location of the LRRK2 gene highlighted with a red underline. f) UCSC Genome Browser view illustrating the genomic location of the VERT region (shaded area) represented in panel e, with the location of the DNAJC6 gene highlighted with a red underline; also highlighted with a green underline is the JAK1 gene. h) UCSC Genome Browser view illustrating the genomic location of the VERT region (shaded area) represented in panel g, with the location of the PRKN gene highlighted with a red underline.

Taken together, these findings underscore the role of I/SCs in regulating disease-relevant gene clusters and emphasizes their conservation, synteny, and functional relevance in both neurological and immune-related disorders. These examples highlight how stochastic epigenetic regulation within I/SCs can contribute to phenotypic variability and disease risk.

## Discussion

The co-occurrence of random monoallelic expression and asynchronous replication between alleles of autosomal genes is well documented ^2–4^, yet its developmental potential and contribution to human disease have not been systematically investigated. In this study, we expand on the discovery and characterization of Inactivation/Stability Centers (I/SCs), by identifying an additional 163 VERT regions using clonal lines of primary ACP cells. This brings the total number of VERT regions to 363, covering approximately 12% of the human genome. In addition, we identified 112 loci where we detected both VERT and AEI, which represent high confidence I/SCs (Supplementary Table 4).

To assess the evolutionary conservation of I/SC regulation, we analyzed replication timing data from mouse pre-B cell clones, identifying 117 mouse VERT loci. Many of these regions overlap with known mouse AEI genes and gene clusters including olfactory, vomeronasal, taste and antigen receptors as well as protocadherin gene families. Notably, we identified 21 syntenic VERT regions shared between human and mouse genomes. These findings reinforce and extend previous observations indicating a high degree of conservation of AEI genes ^4^ and suggest that I/SCs are a fundamental and widespread feature of autosomal epigenetic regulation in mammals.

### I/SC regulation imparts tissue-relevant variation

Our clonal analysis of ACP and LCL cells, combined with haplotype-phased replication-timing and expression profiling, allowed us to identify I/SCs that appear tissue-enriched as well as a subset that is shared across tissues and conserved between species. Although the number of clones and individuals examined here is limited, these observations are consistent with a model in which some I/SCs serve broader or more ubiquitous functions, while others are preferentially engaged in a tissue-dependent manner, reflecting underlying differences in gene expression programs. Consistent with this interpretation, comparisons of VERT regions revealed greater overlap within tissue type (ACP6 vs. ACP7; EB3-2 vs. GM12878) than between tissues. However, given the modest sample size and potential batch effects, these findings should be interpreted as preliminary evidence suggesting, but not definitively establishing, a link between I/SC activity and cellular identity. Expanded analyses across additional tissues, individuals, and clonal lines will be required to rigorously define the extent of tissue specificity in I/SC regulation.

Importantly, we identified 94 VERT regions shared across two or more clonal sets, representing I/SCs that likely reflect core regulatory regions active across individuals and cell types (Figure 1i). As detailed in Supplemental Table 1, we identified VERT regions detected exclusively in single clone sets. These private VERT regions may reflect inter-individual variability in epigenetic regulation, context-dependent I/SC usage, but more likely reflect a limitation of sampling—i.e., regions that would be detected in additional clones if a larger number were analyzed. Thus, while our findings provide strong support for both shared and tissue specific I/SCs, they also underscore the importance of clone number and diversity in defining the full landscape of autosomal epigenetic regulation.

In the context of development, AEI introduces stochastic variability in gene dosage across clonal populations, contributing to cellular heterogeneity during lineage specification and tissue formation. Such heterogeneity could serve as a substrate for cellular selection, enabling the developing tissue to fine-tune function or respond adaptively to environmental cues. This model aligns with observations that AEI often affects genes involved in cell adhesion, signal transduction, and neurodevelopment, where even small variations in expression levels could significantly impact cell fate and connectivity.

At the molecular level, I/SCs contribute to cellular individuality by enabling stochastic monoallelic expression of specific genes. This mechanism is particularly critical in biological systems that require diversity at the single-cell level, such as the olfactory, vomeronasal, immune, and nervous systems. Consistent with this, many I/SCs coincide with gene clusters, including olfactory, vomeronasal, and antigen receptor genes, as well as HLA and protocadherin genes, all of which are known to undergo stochastic monoallelic expression. We identified numerous additional gene clusters, e.g. taste receptors, residing within I/SCs, suggesting that these other gene families may also participate in tissue mosaicism and/or cellular individuality. Our current model proposes that I/SC regulation is integral to tissue development, where stochastic, epigenetically driven allelic regulation introduces phenotypic variability among cells. This variability may promote functional diversity and selective advantage during development. Under this model, each tissue would engage a distinct but partially overlapping set of I/SC-regulated genes, aligned with its unique transcriptional and functional requirements. This framework positions I/SCs as key players in promoting cellular individuality, tissue mosaicism, and developmental plasticity.

Many of the genes that reside within I/SCs are regulated by allelic exclusion. Allelic exclusion is a regulatory mechanism by which only one allele of a gene is expressed while the other is actively silenced, ensuring monoallelic expression in each cell. This process is well characterized in the immune system, where it plays a critical role in B and T cell receptor gene rearrangements, ensuring that each lymphocyte produces a single antigen receptor ^18^. Furthermore, functional monoallelic expression is achieved via a secondary mechanism that involves product inhibition. For example, in developing B cells, expression of a productive immunoglobulin heavy chain allele suppresses rearrangement and expression of the second allele, maintaining clonal specificity ^18^. Allelic exclusion has also been observed in olfactory receptors, where product feedback ensures expression of a single allele of only one member of a large gene family, ensuring that each olfactory receptor neuron responds to a single odorant^2,33,58–61^.

### I/SC regulation acts through mechanisms distinct from X inactivation

Numerous studies in both human and mouse cells support the notion that allele-specific epigenetic regulation of autosomal protein-coding genes is widespread ^4,11,13,37,38,40,58,62,63^. Although stochastic monoallelic expression on autosomes may appear similar to X chromosome inactivation (XCI), the mechanisms behind them are distinct: First, XCI is initiated in preimplantation embryos, as early as the 4-8 cell stage in the mouse and 8-16 cell stage in humans ^64–69^, and later becomes established during differentiation of the inner cell mass of the blastocyst in both species ^68,70^. In contrast, autosomal stochastic monoallelic expression appears later in development, with allelic biases that can be erased and reestablished during differentiation ^10,38^. For example, a recent study found that allele-specific autosomal inactivation is absent during early blood lineage specification but detectable in hematopoietic stem cells and their differentiated progeny ^9^. Second, in XCI, there is strong coordination along the chromosomes: genes on the active X chromosome share both expression and early replication profiles, whereas genes on the inactive X show silencing and late replication. In contrast, autosomal inactivation lacks such coordination: allelic expression and replication timing are unlinked within a chromosome and independent across the genome ^1,7,71^. Taken together, these observations indicate that autosomal allelic inactivation is established through a mechanism that is fundamentally distinct from XCI and arises at a different developmental stage, underscoring that the two processes represent separate epigenetic systems.

### An alternative model for haploinsufficiency

The relationship between genotype and phenotype is fundamental to understanding human genetic disease. At the scale of the whole genome, loss of function alleles are generally not deleterious when heterozygous and are therefore haplosufficient ^21,24^. Not surprisingly, most of the loss of function mutations in single genes in humans are recessive ^21,29^. Therefore, for most human disease genes a single normal allele is sufficient to achieve the typical phenotype. Dominantly inherited disorders comprise a diverse group of conditions that are often severe and poorly understood, except in cases of gain-of-function mutations, where dominant negative or interfering activities induce the abnormal phenotype. However, many dominantly inherited diseases result from copy number changes, either gain or loss of wild-type alleles ^29,72^. Haploinsufficiency, predicted to apply to as much as 40% of the human genome ^19,20^, refers to a heterozygous deletion or complete loss-of-function mutation that results in an altered phenotype, indicating that a single normal allele is insufficient. The underlying molecular mechanisms responsible for most haploinsufficient human disease genes remain largely unknown.

The existence of two distinct populations of cells in XX individuals, one expressing genes from the maternally derived X chromosome and the other from the paternally derived X, has profound implications for both development and the manifestation of X-linked diseases, where both dominant and recessive inheritance patterns are observed in females ^29^. A key factor influencing these inheritance patterns is the presence or absence of cellular selection during development. In certain X-linked disorders, strong selection favors the survival or expansion of cells in which the mutant allele is on the inactive X chromosome, resulting in a predominance of wild-type cells and a clinical presentation consistent with X-linked recessive inheritance ^21,26^. In contrast, when cellular selection is absent, heterozygous females typically retain a ∼50:50 mosaic of wild-type and mutant-expressing cells. The presence of mutant-expressing cells, essentially null cells, is thought to underlie X-linked dominant phenotypes ^27,73^. In such cases, heterozygous loss of function mutations leads to a condition of haploinsufficiency, in which the presence of a single normal allele is insufficient to maintain normal function. Interestingly, although single X-linked alleles are sufficient in males, the mosaic expression pattern in females unmasks phenotypic consequences that are otherwise hidden. This highlights an underappreciated mechanism of haploinsufficiency: the cellular mosaicism inherent to X inactivation can reduce effective gene dosage below functional thresholds in individuals with heterozygous mutations, particularly in tissues where cells expressing the mutant (null) allele persist.

One of the most striking implications of I/SC regulation is its relevance to human genetic disease, particularly those involving haploinsufficiency. Among the 979 genes mapped within human VERT regions, 272 are associated with monogenic diseases, including 87 with autosomal dominant inheritance, many of which are classified as haploinsufficient ^29^. Our findings offer a new mechanistic model for autosomal haploinsufficiency: in the absence of compensatory upregulation and/or cellular selection mechanisms, stochastic allelic inactivation results in or expands a functionally null cell population, even when one allele is wild type. This mirrors the situation observed in X-linked dominant disorders with loss of function mutations, where the absence of cellular selection results in the persistence of null cells. This model expands upon prior theories of haploinsufficiency that assume strict biallelic expression, which attribute dominant phenotypes to factors such as threshold-dependent expression, stoichiometric imbalance in multiprotein complexes, or inability to buffer expression noise ^22–25^. In addition, our model fits nicely with a previous study that modeled stochastic gene expression of disease genes and proposed “haploinsufficiency syndromes might result from an increased susceptibility to stochastic delays of gene initiation or interruptions of gene expression” ^74^. Our data suggest that epigenetically regulated stochastic AEI can contribute to pathogenic phenotypes in heterozygous individuals by either reducing the number of functional cells below a critical disease threshold or by disrupting the normal mosaicism of gene expression within disease-relevant tissues. This expands the conceptual framework of haploinsufficiency to include epigenetic, rather than purely genetic, sources of dosage reduction.

### Limitations of our study

While our findings provide substantial evidence for the existence and functional relevance of I/SCs, several important limitations remain to be addressed. First, the number of clones analyzed in our studies, while sufficient to identify robust patterns of AEI and VERT, limits our ability to capture the full spectrum of stochastic epigenetic states across the genome. For example, we frequently observe AEI of genes located adjacent to VERT regions, indicating that the boundaries of the epigenetic mechanisms functioning at I/SCs remain undefined. This raises important questions about how far the influence of I/SC-associated regulation extends beyond the VERT region. In addition, the specific epigenetic marks, including DNA methylation, histone modifications, and chromatin accessibility, at key regulatory elements such as replication origins, enhancers, and promoters within I/SCs have yet to be defined. Furthermore, the higher-order chromatin organization of I/SCs, including their relationship to topologically associating domains (TADs) and A/B compartments ^49,75^, remains largely unexplored and may hold critical insights into their regulatory architecture and functional constraints. Second, our analysis was conducted in a restricted number of cell types, namely LCLs and ACPs, and therefore does not represent the full diversity of I/SC usage across human tissues. Third, our study was limited to a small number of individuals, which may constrain the identification of inter-individual variability in I/SC regulation. Fourth, the developmental timing of I/SC establishment remains unresolved; it is unclear when during lineage commitment or differentiation these regions acquire their allele-specific states. A related and equally important question is whether I/SCs can be reset during reprogramming into induced pluripotent stem cells (iPSCs) and subsequently reestablished upon differentiation. This issue is particularly relevant given the widespread use of iPSCs in human disease modeling and regenerative medicine ^76–78^. Understanding how I/SCs behave during cellular reprogramming and lineage specification will be critical for interpreting AEI-related phenotypes in iPSC-derived systems.

An additional open question is how the magnitude of stochastic AEI influences disease states, particularly in the context of variable penetrance and expressivity. For example, a recent study employed allelic expression ratios to identify functionally relevant AEI in genes associated with inborn errors of immunity, showing that individuals with the same pathogenic mutation can exhibit divergent clinical outcomes depending on the degree of allelic imbalance ^28^. These findings suggest that AEI can modulate disease severity, acting as an epigenetic layer of regulation that determines whether a given mutation manifests clinically. We anticipate that the biological significance of AEI will vary depending on the function of the gene, its dosage sensitivity, and the cellular context in which it is expressed. Addressing these questions will be essential for achieving a comprehensive understanding of the formation, function, and disease relevance of I/SCs in human biology.

## Conclusion

In conclusion, our work extends the catalog of I/SCs in the human genome and supports a model in which stochastic, allele-specific epigenetic regulation at autosomal loci contributes to cellular diversity and disease susceptibility. Moreover, by identifying syntenic I/SCs in the mouse genome, our findings greatly enhance the potential utility of mouse models for mechanistic studies of disease-causing mutations at conserved loci. Our findings present a potential mechanism for incomplete penetrance, variable expressivity, and tissue-specific vulnerability, particularly in disorders caused by haploinsufficiency. In such cases, stochastic silencing of the wild-type allele in a subset of cells may lead to functional null states, even in heterozygous individuals, reducing the effective number of functional cells below a critical threshold. This may be especially relevant in the nervous system, where precise gene dosage is often essential for maintaining neuronal function. Moreover, I/SC regulation at disease genes raises the possibility that stochastic epigenetic regulation could act as an epigenetic modifier of disease risk, helping to explain why some individuals carrying pathogenic mutations remain asymptomatic. Understanding the full scope of I/SC regulation in both normal development and disease pathogenesis will be essential for identifying novel regulatory mechanisms and potential therapeutic targets. As tools for single-cell and allele-resolved epigenomics continue to advance, I/SCs may emerge as key elements in both basic biology and precision medicine.

## RESOURCE AVAILABILITY

Further information and requests for resources and reagents should be directed to and will be fulfilled by the lead contact, Mathew Thayer.

### Materials availability statement

All data generated or analyzed during this study are included in the manuscript and supporting files. The Repli-seq and RNA-seq raw sequencing data generated in this study have been deposited in the European Nucleotide Archive database under accession code PRJEB110291 and are available without restriction.

## Supporting information

Response to Reviews

## Acknowledgments

M.J.T. was supported by NIH (R01GM130703, and RF1NS142814).

M.B.H was supported by NIH NCI 4K00CA245677-03 DMG was supported by NIH R01GM083337

## Author Contribution Statement

MBH, MJT, and DMG conceived of the experimental design for the RNA expression and replication timing assays. KF performed the surgeries to obtain the amputed digits. BJ and PAY performed the ACP cultures and clone generation. MBH and AEV carried out the RNA-seq assays, including data analysis. MBH and AEV carried out the Repli-seq assays, including data analysis. PFC and PTS provided advice during preparation of the manuscript. MBH, DMG and MJT wrote the manuscript.

## Methods

### Cell culture

All human tissue samples were obtained in accordance with protocols approved by the Oregon Health & Science University Institutional Review Board (OHSU IRB# 00019018) and the Western Institutional Review Board (WIRB# 20193303). Informed consent was obtained from all donors or their legal guardians. ACP cells were grown in Dulbecco’s Modified Eagle’s Medium low glucose (Life Technologies) supplemented with 10% fetal bovine serum (Hyclone). Single cell clones were isolated by plating individual cells in 6 well dishes, and were expanded for >20 population doublings ^79^. All cells were grown in a humidified incubator at 37°C in a 5% carbon dioxide atmosphere.

### Haplotype Phasing of ACP cells

Short-read whole genome sequencing was obtained from two parent/child trios with a range of ∼7-14x genome coverage. BWA ^80^ was used for alignment, and FreeBayes ^81^ used to call putative germline variants jointly for each trio. Variants were filtered to remove sites with Genotype Quality score of less than 30, and sites with missing genotypes for any member of the trio were removed. WhatsHap ^82^ was then used to phase variants in the child utilizing mendelian inheritance patterns and reads containing multiple variants. 1,679,207 high-confidence sites were phased in ACP7, and 837,323 sites were phased in ACP6.

### Allele-specific Repli-Seq

Short read sequence data was aligned with BWA with standard parameters. BAM files were then deduplicated and sorted with SAMtools ^83^. BCFtools mpileup ^83^ was then used to produce allele-specific read counts at phased heterozygous loci. Library size normalization was performed to produce counts per million informative reads. Read counts were then grouped into 250kb sliding genomic windows with no overlap. E/L Repli-Seq values were generated by taking the Log base 2 of early versus late values^84^. Quantile normalization was performed on each group of subclones within each individual sample to allow for comparison among subclones. For each sample, variability of allele-specific RT among subclones was defined as the Standard Deviation (degrees of freedom = 1) of repli-seq at each 250kb window. Genome-wide std. dev. values generated a log-normal distribution, and we set a threshold of Z>=2.25 on the log-normal to identify regions with highly variable allele-specific RT among subclones.

### Allele-specific RNA-seq

Nuclei were isolated by centrifugation for 0.5 minutes from ACP clones following lysis in 0.5% NP40, 140 mM NaCl, 10 mM Tris-HCl (pH 7.4), and 1.5 mM MgCl_2_. Nuclear RNA was isolated using Trizol reagent using the manufacturer’s instructions, followed by DNase treatment to remove possible genomic DNA contamination. Briefly, ribosomal RNAs were removed using the Ribo-Zero kit (Illumina), RNA was fragmented into 250-300bp fragments, and cDNA libraries were prepared using the Directional RNA Library Prep Kit (NEB). Paired end sequencing was done on a NovaSeq 6000 at the OHSU MPSSR core facility.

Strand specific short read sequence data was aligned with STAR ^85^. SAMtools ^83^ was used to remove duplicates and sort BAM files. Strand-specific, allele-specific reads at heterozygous phased SNPs were counted using BCFtools mpileup ^83^. For gene-centric allele-specific analyses, the GENCODE v44 Gene annotation GTF file was used to define gene start and stop coordinates. The informative strand-specific, allele-specific read counts for each heterozygous SNP were then pooled together for all SNPs within each gene. Allelic expression imbalance was calculated as the fraction of reads derived from the maternal vs the paternal allele.

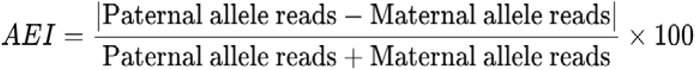

For example, completely balanced allelic expression is 50% AEI, and completely imbalanced AEI is 100%. A threshold of AEI>= 70%, and FDR corrected binomial p-value<=0.05 was used to classify AEI in each individual subclone. The standard deviation (degrees of freedom=1) was used for each gene to generate a metric of AEI variability among subclones. The genome-wide distribution of std. dev. was generated across all genes, and the top 2.5% of variable genes were classified as highly variable among subclones.

**Supplementary Figure 1. Imprinted regions display asynchronous replication.** a and d) VERT regions on human chromosome 14 and 20. The standard deviation in 250 kb windows (circles) are shown. Outlier windows from all four sets of clones (ACP6, ACP7, EB3-2, and GM12878) are highlighted in different colors as shown. The regions highlighted by the red boxes mark known imprinted loci. b and e) For replication timing, solid lines represent the paternal allele (Hap1) and the dotted line represents the maternal allele (Hap2). The regions with asynchrony are shaded and the gene expression is shown for each clone in a different color. c) UCSC Genome Browser view of the asynchronous region in panel a. c and b) The single asterisk marks the paternally expressed DLK gene, and the double asterisks mark the maternally expressed MEG8 gene. f) UCSC Genome Browser view of the VERT region in panel d. f and e) The # marks the paternally expressed L3MBTL1 gene.

**Supplementary Figure 2. VERT detected at the mouse T cell receptor alpha locus.** UCSC Genome Browser view of the mouse VERT region (shaded) detected at the T cell receptor alpha (Tra) locus on mouse chromosome 14. The location of 37 Or genes, marked by red underlines, and the vomeronasal receptor gene Vmn2r88, marked by a green underline.

**Supplementary Figure 3. Mouse syntenic regions display VERT.** a, c and e) VERT regions on mouse chromosome 16, 2, and 18. The standard deviation in 50 kb windows (circles) is shown. Outlier windows from pre-B cell clones are highlighted in orange. The VERT regions highlighted by red boxes are expanded in panels b, d, and f. b) UCSC Genome Browser view of the VERT region in a above, highlighting the location of the mouse App gene (red box). The shaded area represents the VERT region. d) UCSC Genome Browser view of the VERT region in panel c, highlighting the location of the mouse Mocos and Fhod3 genes (red boxes). The shaded area represents the VERT region. f) UCSC Genome Browser view of the VERT region in panel c, highlighting the location of five mouse Scn genes (red boxes). The shaded area represents the VERT region.

## RESOURCE AVAILABILITY

Source data are provided with this paper. Further information and requests for resources and reagents should be directed to and will be fulfilled by the lead contact, Mathew Thayer.

## Materials availability statement

All reagents generated in this study will be made freely available upon request.

## Data Availability Statement

The Repli-seq and RNA-seq raw sequencing data generated in this study have been deposited in the European Nucleotide Archive database under accession code PRJEB110291 and are available without restriction.

