## Supplementary material for "Autosomal Allelic Inactivation: Variable Replication and Dosage Sensitivity": Response to Reviews

**Manuscript number:** RC-2025-03163

**Corresponding author(s):** Mathew Thayer

### 1. General Statements

We thank the reviewers for their thoughtful and constructive comments, which substantially improved our manuscript. In response, we have revised the text and figures throughout to address the points raised. Specifically, we have:

i. Refined our definition of Inactivation/Stability Centers (I/SCs): We limit this designation to loci where both Allelic Expression Imbalance (AEI) and Variable Epigenetic Replication Timing (VERT) were detected, either in the present study or in previously published work.
ii. Expanded methodological clarity: We provide detailed descriptions of how VERT regions were identified, annotated, and quantified, including thresholds for allelic imbalance, replication timing variability, and sampling depth. We also justify the ≥80% AEI cutoff, which is based on recently published studies showing that modest allelic biases can have biological and clinical significance.
iii. Enhanced benchmarking and validation: In addition to the analysis of X inactivation in female ACP cells, we now include comparisons between imprinted and non-imprinted regions to benchmark the magnitude of allelic replication timing imbalance, demonstrating that the magnitude of imbalance observed at non-imprinted VERT regions is comparable to known imprinted regions.
iv. Address tissue specificity and sampling limitations: We now discuss how the data derived from a limited number of clones, tissues, and individuals support the identification of robust AEI and VERT patterns. In the future, additional tissues and individuals will be required to capture the full diversity of I/SC regulation.
v. Clarify biological relevance: We have expanded our discussion to highlight the consistency of AEI findings across cell types, including examples of genes implicated in neurodevelopmental and neurodegenerative disorders, and we clarify our model of how I/SC regulation contributes to haploinsufficiency, variable expressivity, and incomplete penetrance in human disease.
vi. Improved figures and supplemental data: We have updated figure legends for clarity, added a new supplementary figure benchmarking imprinted regions, added supplementary tables containing: the full description of our GO analysis, the list of I/SCs where we have detected both VERT and AEI, the ratios of the number of transcripts derived from early and late replicating alleles for the I/SCs illustrated in all figures, and we have cross-referenced all supplementary tables.

### 2. Point-by-point description of the revisions

**Reviewer 1:** “The existence of VERT regions is well supported, but the number of regions called as ISCs may be inflated by permissive thresholds (e.g., AEI {greater than or equal to} 0.8 or {less than or equal to} 0.2 in a single clone). This risks conflating transient stochastic differences with stable ISCs.”

**Response:** We selected the ≥80% (or ≤20%) allelic imbalance threshold, along with the requirement of at least one biallelic clone, as our criterion for significant AEI. This choice was guided by a recent study demonstrating that allelic imbalance, as low as a 65%/35%, is enough to effect disease penetrance in humans (Nature 2025; 637:1186–1197). For completeness, results obtained using more stringent thresholds (≥90% and ≥95% imbalance) are presented in Supplementary Table 2.

Furthermore, it is unlikely that transient stochastic differences in allelic expression, such as those detected by single-cell RNA sequencing assays (Nat. Rev. Genet. 2015; 16:653–664), would be captured by our approach. Each clone in our study was expanded from a single cell to over one million cells before both RNA-seq and Repli-seq analyses, effectively averaging out transient transcriptional and/or replication fluctuations, and thus reflecting stable, mitotically heritable epigenetic states.

**Reviewer 1:** “More robust approaches would include using magnitude of imbalance, annotating VERTs by genomic location, applying stricter thresholds for replication timing, and benchmarking AEI distributions against the X chromosome.”

**Response:** All VERT regions identified in this study were annotated according to both the magnitude of allelic imbalance and their genomic coordinates, using 250 kb windows for the human samples and 50 kb windows for the mouse samples (see Supplementary Tables 1 and 6). Figure 1c directly compares the magnitude of imbalance, defined as outliers in the standard deviation, for both allelic replication timing and allelic expression across autosomal and X-linked loci in female ACP cells.

In addition, we detected allelic replication asynchrony at 12 known imprinted loci, and the standard deviation of replication timing at these loci, measured in 250 kb windows, is comparable to that observed across the >350 VERT regions detected at non-imprinted sites. For comparisons, we have highlighted the imprinted regions with + symbols in Figures 1e, 2d, 3c, 6g, 7e, 7g, and we have highlighted the imprinted regions in Supplemental Table 1, and in the Data Source files. For additional comparisons, we have included Supplemental Figure 1 to illustrate the magnitude of replication timing imbalance and allele-specific gene expression at two autosomal imprinted regions.

**Reviewer 1:** “Figures and text would benefit from improved clarity: axis labels are missing in places (e.g., Fig. 1c, Fig. 2g), legends should explain chromosome arm colors, and cluttered figures such as Fig. 1j could be re-visualized for interpretability.”

**Response:** Figure labels have been added to Figs. 1c and 2g, and legends modified for clarity.

**Reviewer 1:** “the claim of cell-type specificity is not convincingly demonstrated given the small sample size (n=4) and strong batch confounding between lymphoblastoid and cartilage progenitors.” And “Hierarchical clustering is confounded by batch and based on presence/absence calls that lack quantitative resolution.”

**Response:** We agree that the limited number of individuals and clones, as well as the comparison between only two distinct tissue types (LCLs and ACPs), have quantitative limitations. Our primary intent was to evaluate whether any I/SCs were shared between independently derived clonal cell lines from different tissues to determine whether there is evidence of tissue-specific I/SC usage, rather than to make quantitative claims about global cell-type specificity.

To address this concern, we have replaced the hierarchical clustering analysis, in Figure 1i, with a Venn diagram that more directly illustrates the overlap and tissue-specific distribution of VERT regions detected in the different clonal sets. This revised representation avoids assumptions about clustering relationships and removes batch-driven bias, while still conveying the key observation that many VERT regions are shared across tissues and others appear tissue-restricted.

**Reviewer 1:** “While syntenic VERT regions across mouse and human are intriguing, they complicate interpretation of strong clustering by cell type. Sampling depth may also have exaggerated allelic imbalance calls.”

**Response:** We note that the human LCLs used in our study are B cells, and immunoglobulin gene rearrangements were used to confirm the clonal uniqueness of each line. Similarly, the mouse replication timing data analyzed here was generated from pre-B cells, which also undergo immunoglobulin gene rearrangements. Thus, both the human LCL and mouse pre-B cell datasets were derived from B-cell lineages, providing a consistent cellular context for comparative analysis.

Sequencing depth is an important consideration for all variant base calls. Without fully haplotype-resolved genomes, previous studies relied on calculating per-SNP calls of allelic imbalance based on reads covering a single nucleotide locus. To improve sequencing depth supporting the identification of VERT and AEI regions, we utilized haplotype-resolved genomes that allowed all informative allele-specific reads to be pooled across all heterozygous SNPs within genomic windows or expressed genes. For AEI, we set a minimum threshold of 20 informative allele-specific reads per gene, a minimum FDR-corrected p-value of <=0.05, and a minimum of 80% vs 20% allelic imbalance. Importantly, a recent study showed that allelic imbalance as low as a 65%/35% is clinically relevant in humans (Nature 2025; 637:1186–1197). We reiterate that more stringent thresholds (≥90% and ≥95% imbalance) are presented in Supplementary Table 2.

**Reviewer 1:** “Gene set enrichment analysis should be restricted to avoid inflated significance from overly broad categories.”

**Reviewer 2:** “Some of the GO terms presented are too broad to suggest any biological significance to the result, even if there is statistical significance (for example, the top term for LCL clones 'Cytoplasm' is associated with 12,000 genes, and the second term for mouse clones 'Membrane' is associated with 10,000). It would be helpful to focus on GO terms lower in the GO hierarchy.”

**Response:** We now include our complete Gene Ontology analysis, with more specific biological categories, in Supplemental Table 5.

**Reviewer 2:** “Allelic imbalance has been referred to as AI, MAE (monoallelic expression), RMAE (random monoallelic expression) etc. The paper whose mouse data the authors make use of uses Asynchronous Stochastic Replication Timing (ASRT) instead of VERT to refer to the same phenomenon. Creating unnecessary jargon makes the paper more difficult to read and adds needless complexity to an already complex field.”

**Response:** While we agree that allelic expression imbalance has been described by different investigators using many different phrases, we believe that MAE, RMAE and AI do not represent an accurate description of the phenomenon. In our study [and our previous study; Nat Commun. 2022; 13(1):6301] we used clonal analysis of allele-specific expression and found that while some clones display equivalent levels of expression between alleles of a given gene (i.e. bi-allelic expression) other clones express only one allele (i.e. mono-allelic expression), and yet other clones have undetectable expression (i.e. silent on both alleles). This pattern of allele-restricted expression indicates that each allele independently adopts either an expressed or silent state. Importantly, because these expression states are mitotically stable, allele-autonomous, and independent of parental origin, we refer to the choice of the expressed allele as stochastic. Given this variability, we believe that the phrase “Allelic Expression Imbalance” (AEI) represents a more accurate descriptor for this phenomenon. We also point out that “Allelic Expression Imbalance” has also been used by other investigators >120 times in the Pubmed database.

In addition, the replication asynchrony that exists at these loci is not consistent with purely ASynchronous Replication Timing (ASRT) between alleles. We found that each allele can independently adopt either earlier or later replication timing in different clones. This variability results in some clones exhibiting pronounced asynchrony between alleles, while in others, the two alleles replicate synchronously, with both adopting either the earlier or later timing state. As reported in our previous study (Nat. Commun. 2022; 13:6301), this behavior reflects a stochastic and allele-autonomous process, leading us to describe these loci as exhibiting Variable Epigenetic Replication Timing (VERT), which we believe is a more accurate descriptor of this phenomenon.

**Reviewer 2:** “The point that allelic imbalance is enriched in VERTs would be enhanced if the authors could present the allelic ratio for all genes found in all VERTs, demonstrating how replication timing on either chromosome affects the allelic ratio.”

**Response:** The stochastic nature of allelic expression and replication timing observed at VERT loci indicates that each allele independently acquires its epigenetic state. In addition, there are typically more than one transcription unit, both protein coding and non-coding, within each VERT region, and each transcription unit also acquires its expressed or silent state independently. Therefore, the expressed or silent status of one allele of a transcription unit does not predict the replication timing or expression status of the same or opposite allele of any other transcription unit within the VERT region. Accordingly, the Early/Late pattern of replication timing that we detect, both in this study and in our previous work (Nat. Commun. 2022; 13:6301), is not correlated with which allele is transcriptionally active. This supports our conclusion that asynchronous replication timing is not a downstream consequence of monoallelic transcription, but rather an independent epigenetic feature of I/SCs. Regardless, because each transcription unit is independent, we provide the expression ratios for all transcripts that are generated from the VERT regions for the coding and non-coding transcription units in Figures 1, 2, and 6; shown in Supplemental Table 9. This analysis indicated that 4,017 informative reads were derived from the earlier replication allele and 3,161 informative reads were derived from the later replication allele, generating an allelic ratio of 1.3 (early/late) and a binomial P value of 1.0.

In addition, a similar analysis of imprinted loci reveals that even at genomic regions with parent-of-origin–specific expression, the replication timing of each allele does not align with transcriptional activity, i.e. both early- and late-replicating alleles can be transcriptionally active, depending on the gene. This observation is consistent with the complex organization of many imprinted domains, where genes on opposite alleles exhibit reciprocal expression patterns. To illustrate this point, we now include Supplemental Figure 1 demonstrating that imprinted loci harbor genes expressed from both the earlier- and later-replicating alleles. In addition, quantification of the total number of transcripts at the DLK1/MEG8 imprinted locus (Supplementary Figure 1a-1c) indicates that the ratio of transcripts derived from the early versus late replicating alleles is equivalent (i.e. a ratio of 1.0; See Supplemental Table 9).

**Reviewer 2:** “Figure 3 highlights the association of related gene clusters with VERTs but the VERTs are assigned based on variable replication timing in just 1 or 2 clones. This is an interesting observation, but to make the point that "VERT regions frequently coincide with gene clusters in the human genome" there needs to be a systematic assessment of replication timing at all gene clusters across all clones, and a statistical test for significance.”

**Response:** Our intent in Figure 3 was not to suggest that all gene clusters are subject to VERT and AEI, but rather to highlight that several well-characterized multigene families that are known to exhibit AEI, such as olfactory receptor, protocadherin, and HLA gene clusters, coincide with VERT regions at their genomic locations. These examples serve as representative illustrations demonstrating that I/SC-associated regulation occurs at established AEI loci organized in gene clusters.

To clarify this point, we have revised the text to explicitly state that Figure 3 presents illustrative examples of known AEI-associated gene clusters overlapping with VERT regions, rather than a comprehensive or statistically exhaustive analysis of all gene clusters across the genome.

**Reviewer 2:** “It is an interesting hypothesis that VERTs are conserved between species at synentic loci. If such regions are really conserved, one would expect that replication timing at these sites would be consistently asynchronous. However the data presented shows that in human clones these VERTs can be specific to an individual donor (as in 5A) or an individual clone (as in 5H).”

**Response:** As discussed in our Limitations Section, our analysis was restricted to a limited number of cell types, clones, and individuals, which may not capture the full diversity of I/SC usage across tissues and populations. While our dataset was sufficient to identify robust patterns of AEI and VERT, it likely represents only a subset of the broader landscape of I/SC regulation in both humans and mice. We anticipate that future studies incorporating a wider range of tissues, individuals, and clonal analyses will uncover an even greater degree of conservation and diversity in I/SC usage across genomes.

**Reviewer 2:** “In order to support the claim that neurodevelopmental disease associated genes reside in asynchronously replicating regions, and are thus more prone to allelic imbalance, the authors would need to demonstrate this phenomenon in neuronal cells.”

**Response:** We make two points that address this critique: First, many of the neurodevelopmental disease genes located within or adjacent to VERT regions are not exclusively expressed in neuronal cells and have previously been shown to exhibit AEI in non-neuronal contexts. For example, Gimelbrant and Chess (Science, 2007; 318:1136–1140) demonstrated AEI of the Parkinson disease genes SNCA and LRRK2 in lymphoblastoid cell lines (LCLs), and in our previous study, we detected AEI of DNAJC6, another Parkinson disease gene, also in LCL cells (Nat. Commun. 2022; 13:6301). In the present study, using cartilage progenitor cells, we identified VERT and AEI of several epilepsy-associated genes, including SCN1A, SCN2A (Fig. 6b), GABRA1(Fig. 6e), and SAMD12 (Fig. 6j), as well as a gene implicated in autism and neurodevelopmental disorders, SEMA5A (Fig. 5c), indicating that these genes are not exclusive to neuronal cell types.

Second, independent studies from the Dr. E. Heard laboratory have provided further evidence that AEI occurs in neuronal lineages. Using mouse neural progenitor cells (NPCs), they identified genes subject to AEI (Dev. Cell, 2014; 28:366–380) and they later evaluated AEI of syntenic human neurodevelopmental disease genes, including Snca, App, Eya4, and Grik2 (Nat. Commun. 2021; 12:5330). In addition, and consistent with our use of AEI, they used the phrase “Allelic Expression Imbalance” to describe the epigenetic expression biases at these genes.

Together, these findings reinforce that AEI, and by extension I/SC regulation, is not restricted to specific cell types, but rather represents a generalizable mechanism of stochastic epigenetic regulation that includes genes relevant to neurodevelopment and disease.

**Reviewer 2:** “However, the authors consistently lean on thin evidence (i.e. a single clone) within a modestly sized dataset (4 clones from 2 donors each) to propose a new model for haploinsufficiency in human disease. The consistent focus on limited elements in the data and perhaps an overreach in the interpretation makes it difficult to appreciate what is in fact a very good experiment.”

**Response:**

We agree that our analysis was conducted on a modest number of clones and individuals, which we explicitly acknowledge as a limitation of the present study. However, several key points support the robustness and broader relevance of our conclusions:

i. Clonal Design and Replication: The strength of our approach lies in its clonal resolution. Each clone represents a single-cell–derived population expanded to over a million cells, enabling direct detection of stable, mitotically heritable allele-specific epigenetic states that would not be apparent in population-averaged data. Importantly, many of the VERT regions we identified are shared between independent clones from different donors and across distinct cell types (ACP and LCL), demonstrating reproducibility and biological consistency.
ii. Cross-Species Validation: We further identified syntenic VERT regions in mouse pre-B cell clones, including at loci known to exhibit AEI in prior studies, providing independent validation and evolutionary conservation of the phenomenon.
iii. Integration with Published Evidence: Our findings extend prior observations of AEI and VERT (e.g. Gimelbrant et al. Science 2007; Heskett et al. Nat. Commun. 2022) and are fully consistent with known stochastic allelic expression imbalance of autosomal genes. We also draw parallels with the absence of cellular selection mechanisms that dictate dominant inheritance patterns for loss of function alleles for X linked disease genes (reviewed in: J Clin Invest, 2008, 20-23; and Nat Rev Genet. 2025, 26, 571–580). Our proposed model linking I/SC regulation to haploinsufficiency is therefore a synthesis of our results with an extensive body of published data, not an inference drawn from isolated observations.
iv. Scope and Framing: We have revised the manuscript to clarify that our proposed model represents a mechanistic framework, not a definitive or exclusive explanation, for how stochastic allelic regulation could contribute to dosage-sensitive disease phenotypes. We also explicitly discuss the need for larger datasets and additional tissues to refine and test this model.

In summary, while we recognize the limited sampling depth inherent to clonal analyses, the consistency of our observations across donors, cell types, and species, together with prior corroborating studies, supports the validity of the conclusions and justifies the broader conceptual implications.
