## Supplemental Figures for "Autosomal Allelic Inactivation: Variable Replication and Dosage Sensitivity"

**Supplementary Figures.**

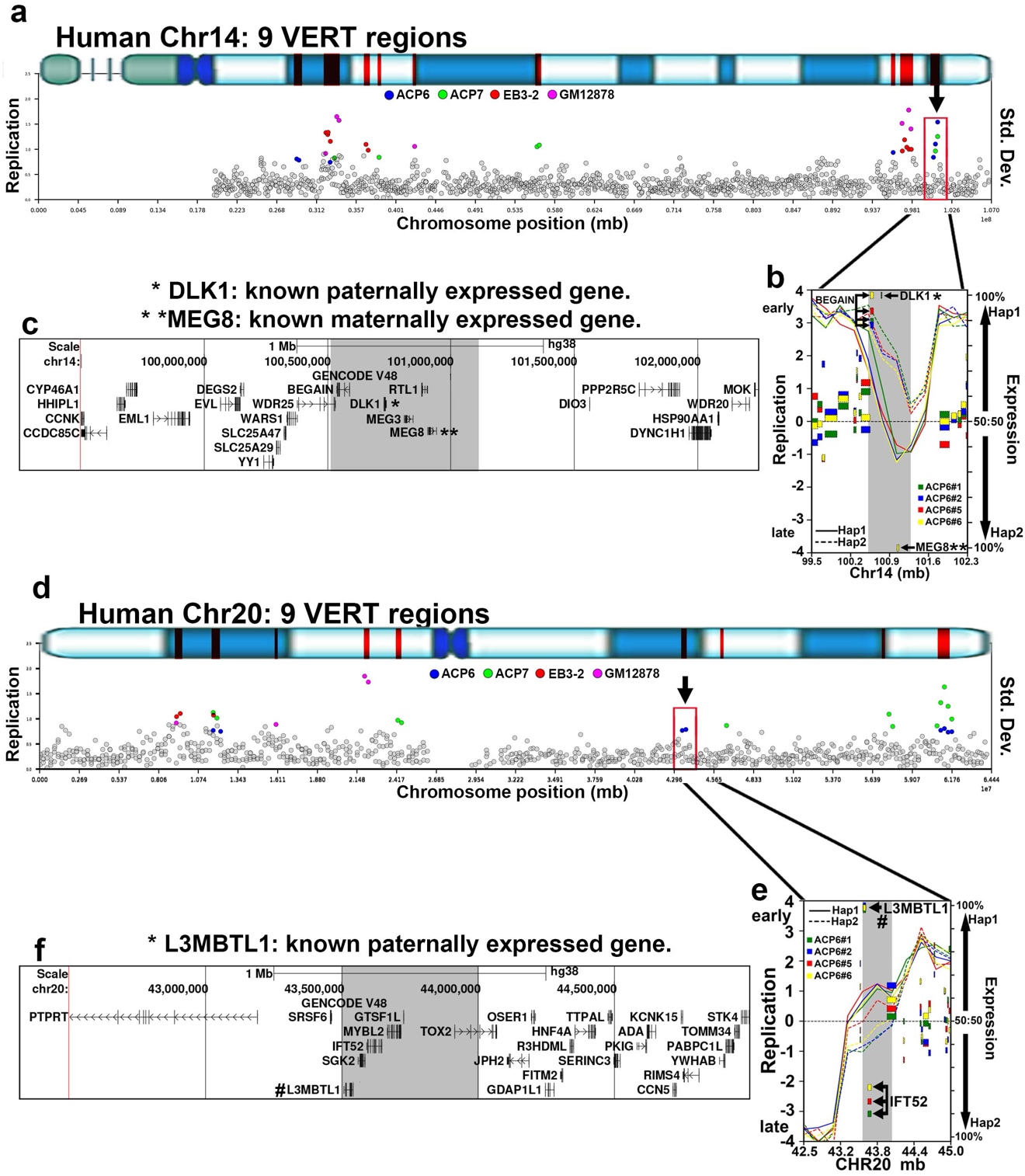

**Supplementary Figure 1. Imprinted regions display asynchronous replication. a and d) VERT regions on human chromosome 14 and 20. The standard deviation in 250 kb windows (circles) are shown. Outlier windows from all four sets of clones (ACP6, ACP7, EB3-2, and GM12878) are highlighted in different colors as shown. The regions highlighted by the red boxes mark known imprinted loci. b and e) For replication timing, solid lines represent the maternal allele (Hap2) and the dotted line represents the paternal allele (Hap1). The regions with asynchrony are shaded and the gene expression is shown for each clone in a different color. c) UCSC Genome Browser view of the asynchronous region in panel a. c and b) The single asterisk marks the paternally expressed DLK gene, and the double asterisks mark the maternally expressed MEG8 gene. f) UCSC Genome Browser view of the VERT region in panel d. f and e) The # marks the paternally expressed L3MBTL1 gene.**

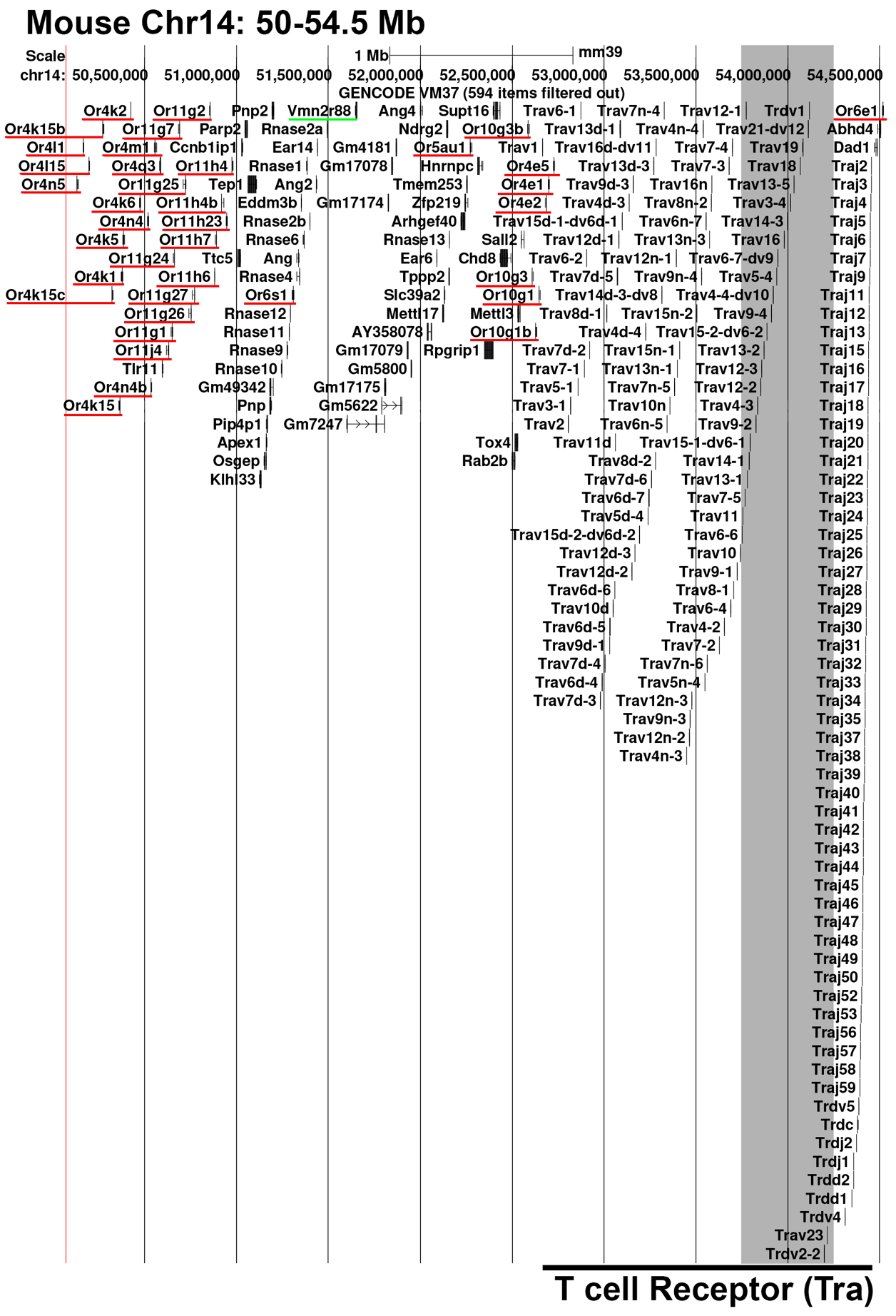

**Supplementary Figure 2. VERT detected at the mouse T cell receptor alpha locus. UCSC Genome Browser view of the mouse VERT region (shaded) detected at the T cell receptor alpha (Tra) locus on mouse chromosome 14. The location of 37 Or genes, marked by red lines, and the vomeronasal receptor gene Vmn2r88, marked by a green line.**

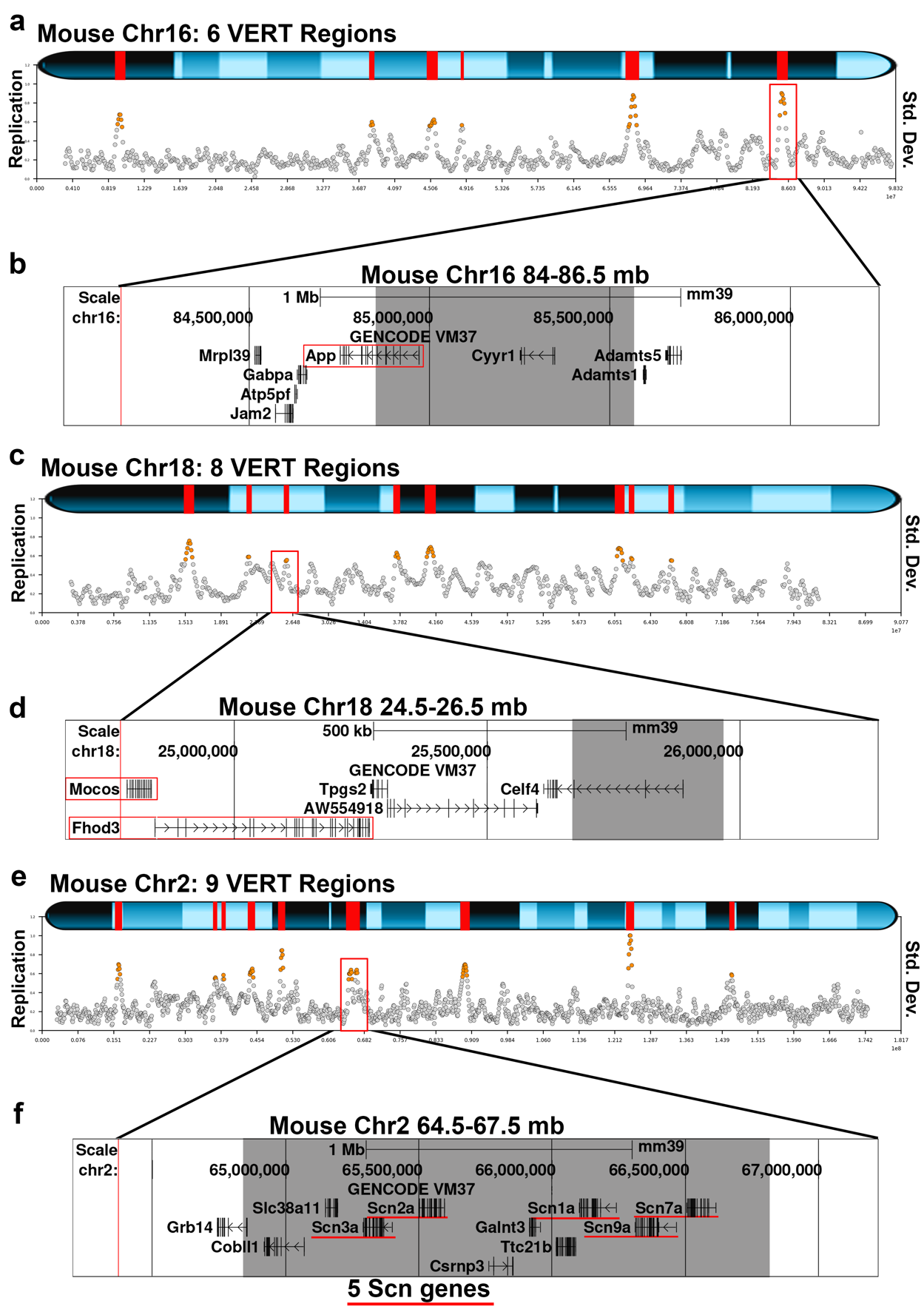

**Supplementary Figure 3. Mouse syntenic regions display VERT. a, c and e) VERT regions on mouse chromosome 16, 2, and 18. The standard deviation in 50 kb windows (circles) is shown. Outlier windows from pre-B cell clones are highlighted in orange. The VERT regions highlighted by red boxes are expanded in panels b, d, and f. b) UCSC Genome Browser view of the VERT region in a above, highlighting the location of the mouse App gene (red box). The shaded area represents the VERT region. d) UCSC Genome Browser view of the VERT region in panel c, highlighting the location of the mouse Mocos and Fhod3 genes (red boxes). The shaded area represents the VERT region. f) UCSC Genome Browser view of the VERT region in panel c, highlighting the location of five mouse Scn genes (red boxes). The shaded area represents the VERT region.**
